# Response of the Oligo-Miocene Bivalve Fauna of the Kutch Basin (Western India) to Regional Tectonic Events

**DOI:** 10.1101/2021.09.08.451389

**Authors:** Saurav Dutta, Devapriya Chattopadhyay

## Abstract

Tectonic changes has influenced the evolution of the marine community by changing the land and seaway configuration through time. Two such tectonic events during Oligo-Miocene times — the closure of the Tethyan seaway due to development of the *Gomphotherium*-Landbridge leading to separation of the Arabian Sea from proto-Mediterranean Sea (∼19 Ma) and significant uplift of the Tibetan plateu marking the initiation of the monsoon (∼16 Ma) — represent a classic case of tectonic shift influencing the regional environment of the Indian subcontinent. We investigated the taxonomic and body size related response of the shallow marine fauna to this regional change using bivalves from 11 time-constrained shellbeds of the Kutch Basin (western India) from three formations — Maniyara Fort (Chattian), Khari Nadi (Aquitanian) and Chhasra (Burdigalian-Langian) representing a time span of ∼9 Ma (24.4 – 15 Ma).

Our collection of over 2000 individuals represents a total of 15 families and 61 morphospecies. The fossils are predominantly calcitic in nature and families of aragonitic composition are often preserved as molds indicating a potential negative effect of diagenesis. The taphonomic nature, however, does not vary substantially across shellbeds and hence, less likely produced a temporal pattern. The five most abundant species, *Ostrea latimarginata, Ostrea angulata, Talochlamys articulata, Anomia primaeva* and *Placuna lamellata* occur in all the formations. The species composition of the Maniyara Fort formation is substantially different from those of the younger formations implying the possible effect of biogeographic separation. Moreover, the absence of proto-Mediterranean taxa in Oligocene shellbeds support a limited faunal exchange as early as ∼24.4Ma (Chattian) ago. We observed a monotonic increase in the overall rarefied species richness and a decrease in evenness from the Maniyara Fort to the Chhasra Formation. However, shellbed analyses show a dominantly conservative behavior of diversity and body size without a strong directional trend through time. Although it is difficult to rule out the negative influence of taphonomy on the diversity of the studied fauna, Oligo-Miocene marine bivalve fauna of the Kutch Basin demonstrate little or no influence of the Tethyan closure and Himalayan upliftment.

## INTRODUCTION

Tectonic shifts often lead to environmental changes, which can drive organisms in a community to change their geographic distribution tracking their preferred environment, or when staying to adapt to the changing environmental conditions (Valentine, 1961, Fields et al, 1993, Roy et al., 1995, Parmesan, 2006, Diniz-Filho and Bini, 2008). Documenting such events in the geologic past is important to understand the nature of faunal response to large-scale environmental changes – which, although crucial in the context of the present environmental crisis, is often impossible to document with short-term historic records (Vitousek et al, 1997). The configuration of the Tethyan seaway represents a classic case of tectonic shift that also influenced regional climate, consequently affecting the distribution of both marine and terrestrial faunas (Renema et al, 2008; Harzhauser et al, 2007).

At the beginning of the Mesozoic, the Neo-Tethys existed as a narrow ocean basin in the east of Gondwana, which subsequently widened (Sengör 1984). The configuration of the Mesozoic Tethys Ocean ended with the dispersal of the Pangean continent and the northward movement of India (Rögl 1999). During the Cenozoic, the Tethyan realm was composed of the western Indo-Pacific Ocean, the Proto-Mediterranean Sea and Paratethys Sea (e.g. Rögl 1998; Popov et al. 2004). During Oligocene and early Miocene times, the Tethys connected two major oceanic areas, the Atlantic and the Pacific oceans. The shallow marine fauna of this time imterval documents two major biogeographic compartments, namely the western Tethys Region (WTR) and the Proto-Indo-West Pacific Region (PIWPR) (Harzhauser et al. 2007). The WTR was composed of the Mediterranean-Iranian Province (MIP) in its center, the eastern Atlantic Province (EAP) in the west and two provinces, the western Indian Province (WIP) and the eastern African-Arabian Province (EAAP), in the east. The WIP consists of the marine faunas of Pakistan and N and SW India. During the early Miocene, a terrestrial corridor called the “*Gomphotherium* Landbridge” developed as a result of the collision between the Afro-Arabian plates and Eurasia (Rögl, 1998, 1999) (Fig. 1). With the rise of this land connection during the early Burdigalian (∼19Ma), the eastern part of Paratethys became disconnected and the seaway between the Mediterranean Sea and the Indian Ocean was closed. Along with changes in the seaway configuration, the early Miocene also witnessed the rise of the Himalayas, which eventually initiated the monsoon – a major climatic event of regional importance. Although the exact timing of monsoon initiation is strongly debated (summarized by Clift and Webb, 2019), the sedimentological record of the western part of the Indian subcontinent constrains it within Burdigalian (Clift et al, 2008 a, b).

**FIG 1.**
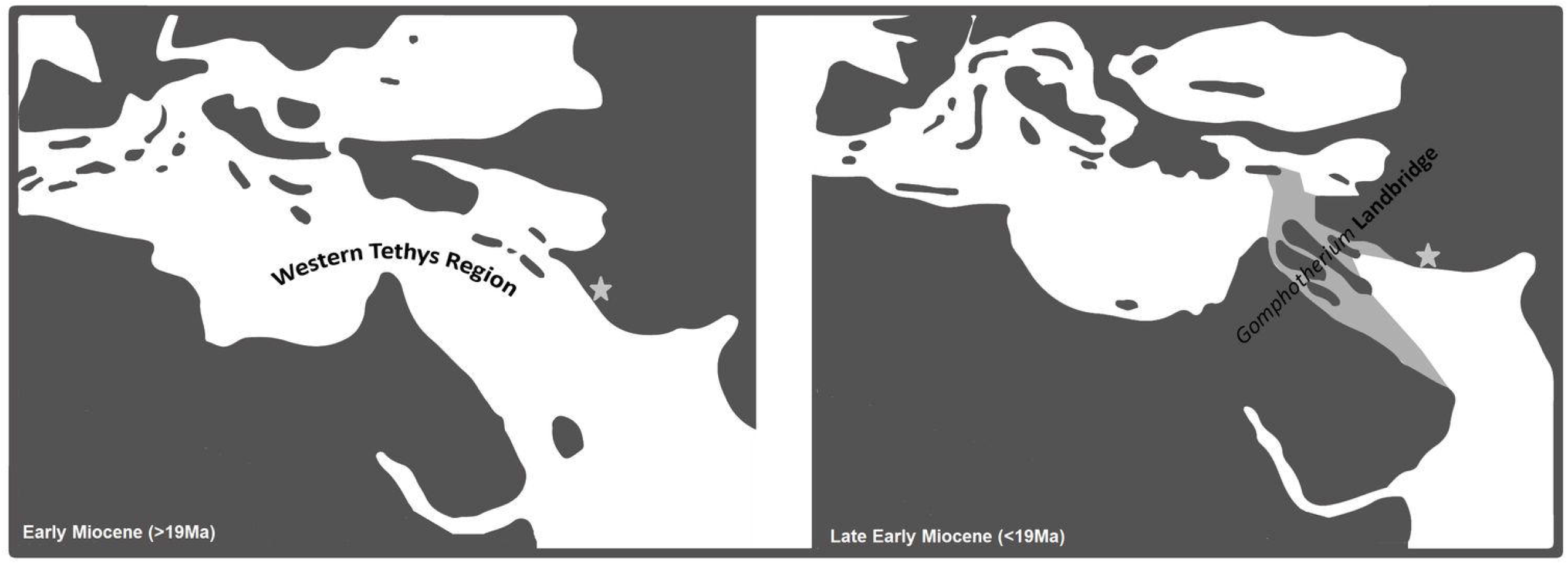
Paleogeographic map of the circum-Mediterranean region before and after the formation of *Gomphotherium* land bridge during the Early Miocene (after Dutta et al, 2020). The star denotes the studied locality from the Kutch basin.

A seaway closure affects the distribution of the fauna, both terrestrial and marine. A change in the regional environment such as the one caused by tectonic uplift influences a variety of physical parameters (such as sedimentation rate, river influx, salinity etc) that can affect the diversity and morphology of shallow marine ecosystems (Scavia et al., 2002, Bakun, 2010, Gilbert et al., 2010, Keeling et al., 2010). Shallow marine communities are more likely to respond to such changes than their deep water counterparts. Although the Oligo-Miocene changes in the Tethyan seaway and Himalayan orogeny has been recognized to play an important role in shaping the terrestrial faunal diversity of the Indian subcontinent (Sahni and Mitra, 1980a, b), the response of shallow marine faunas to such changes from this region is largely understudied with the exception of few broad biogeographic comparisons of shallow marine faunas across the Tethyan seaway (summarized by Harzhauser et al, 2007).

The Late Oligocene-early Miocene deposits of the Kutch Basin in western India preserves a nearly continuous shallow marine succession from the late Oligocene to Miocene. Numerous shellbeds with well constrained age from three formations — Maniyara Fort (Bermoti member – Chattian), Khari Nadi (Aquitanian) and Chhasra (Burdigalian - Langhian) spanning ∼ 9My (24.4 - 15Ma) (Dutta et al, 2020) records deposition during times of some major regional events including (1) closure of the Tethys due to formation of the *Gomphotherium* Landbridge at ∼19Ma (Rogl, 1998, 1999), (2) onset of monsoon intensification ∼16Ma (Clift and Gaeedicke, 2002). In this work, we utilize the fossil record of the diverse bivalve fauna of the Kutch Basin spanning over this crucial time period to evaluate the taxonomic and body size related response to regional tectonic shifts reconfiguring the seaway and influencing the climate.

## MATERIAL AND METHODS

### Study area and geologic setting

The Kutch Basin in western India (Fig. 2A) is a peri-cratonic rift basin that preserves a sedimentary succession from the Triassic to Recent (Biswas, 1992). Tertiary rocks are exposed in the western part of the basin and are dominated by Neogene successions. Oligocene-early Miocene lithounits of the Kutch Basin are primarily of partly carbonate and partly siliciclastic composition and the carbonate units contain diverse assemblages of skeletal invertebrates. Based on sedimentological and microfossil attributes, these units have been interpreted to represent shallow marine settings (Catuneanu and Dave, 2017). We followed a transect along the river Khari and one of its tributaries, river Mithi, where we identified 11 distinct shell beds, two from Upper Maniyara Fort (Bermoti member), one from Khari Nadi and eight from the Chhasra Formation (Fig. 2B). Five of these shellbeds are dated using Sr-stratigraphy (discussed in details by Dutta et al, 2020).

**FIG 2.**
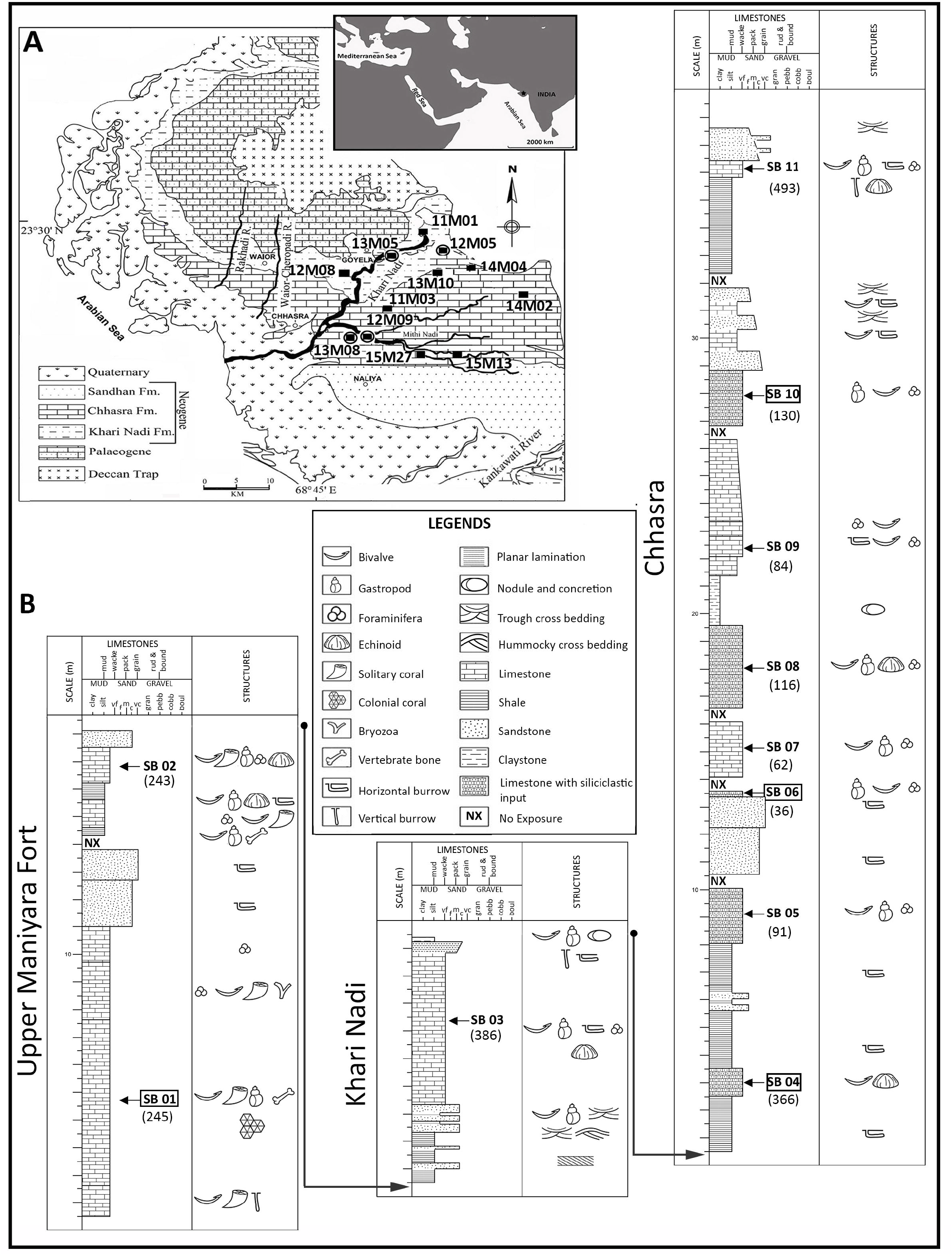
A) Geological map of the studied locality with the map of Indian subcontinent (inset) (modified after Dutta et al, 2020). The solid rectangles represent collection sites of bivalve fossils and the open circles around those rectangles represent the location of the shellbeds which are dated. B) Composite litholog of the three formations (Maniyara Fort, Khari Nadi and Chhasra) from the Oligocene and early Miocene deposits of Kutch (modified after Dutta et al, 2020). The dated shellbeds are represented by a square. The numbers within parenthesis denote the number of bivalve individuals used for analyses.

The extensive presence of corals and complete absence of siliciclastic input in the two shellbeds (SB 01 and SB 02) of Maniyara Fort formation are indicative of the development of coral bioherms that supported a diverse fauna (Table 1, Fig. 3). The Khari Nadi Formation consists of bluish-gray claystone and medium-grained sandstone at the base followed by successions of fossiliferous limestone and shale units. The single shellbed (SB 03) from this formation has a wide lateral extent and is exposed at multiple locations (Dutta et al, 2020). This shellbed is primarily composed of limestone with high siliciclastic input and is rich in fauna (Table 1). We also observe networks of horizontal burrows (Fig. 4). This shellbed is overlain by an intensely bioturbated zone; the burrows are often filled with foraminifera rich sediments. The overlying Chhasra shellbeds are rich in well preserved fauna (Tabe 1, Fig. 5). A clear disconformity is observed on top of SB 11 shellbed, after which the sandy Sandhan Formation starts (Fig. 6F in Dutta et al, 2020).

**Table 1.** A brief description of the sedimentological and paleontological characteristics of the shellsbeds considered in the present study.

| Formation | Shellbed | Description |  |
| --- | --- | --- | --- |
|  |  | Faunal composition | Sedimentology and preservation |
| Chhasra | SB 11 | Gastropods, bivalves, irregular echinoids, barnacles and crustaceans. | Hard-compacted limestone. It is heavily bioturbated with horizontal and vertical burrows. |
|  | SB 10 | Gastropods, bivalves and foraminifera. | Hard-compacted mixed siliciclastic-carbonate. Some of the shells are preserved as molds. |
|  | SB 09 | Echinoderms, foraminifera and bivalves, gastropods. | Loosely-compacted mixed siliciclastic-carbonate with sparsely spread fossil occurrences. |
|  | SB 08 | Bivalves, regular and irregular echinoids, barnacles, gastropods, shark teeth, vertebrate bones and annelid tubes. | Relatively fine-grained limestone. All the bioclasts are preserved in pristine condition. |
|  | SB 07 | Bivalves, barnacles and gastropods. | Loosely-compacted mixed siliciclastic-carbonate with well-preserved shells. |
|  | SB 06 | Bivalves, irregular echinoids, barnacles, foraminifera, bryozoa, sponges and gastropods. | Hard-compacted limestone. Bivalve shells are preserved in their live position and show low degree of disarticulation. Gastropod shells are often preserved as molds. |
|  | SB 05 | Bivalves, echinoids, barnacles, foraminifera and gastropods. | Hard and compact limestone with low degree of disarticulation of the bivalve shells. |
|  | SB 04 | Bivalve, echinoids, barnacles, foraminifera and gastropods. | Fine grained limestone with low degree of disarticulation of the bivalve shells. Thin shelled *Placuna lamellata*, are oriented parallel to bedding and are exceptionally well preserved. |
| Khari Nadi | SB 03 | Irregular echinoids, corals, bivalves, barnacles and gastropods (primarily *Turritella*). | Limestone with high siliciclastic input. Cross stratification is present. Networks of horizontal burrows present. |
| Manyara Fort | SB 02 | Bivalves, gastropods, crab claws, corals, foraminifera and shark teeth. | Fine grained limestone. The bioclasts of the shell beds are coarse and densely packed. The shells are preserved in situ in their pristine form with low to moderate disarticulation, dissolution, fragmentation, pitting and bioinfestation. Internal molds were also abundant in this shellbed. |
|  | SB 01 | Bivalves, gastropods, echinoids, barnacles, foraminifera, bryozoans, vertebrate bones, and corals. | Fine grained limestone. The bioclasts of the shell beds are randomly oriented and the shells are preserved in situ with low to moderate disarticulation, dissolution, fragmentation, pitting and bioinfestation. |

**Fig 3.**
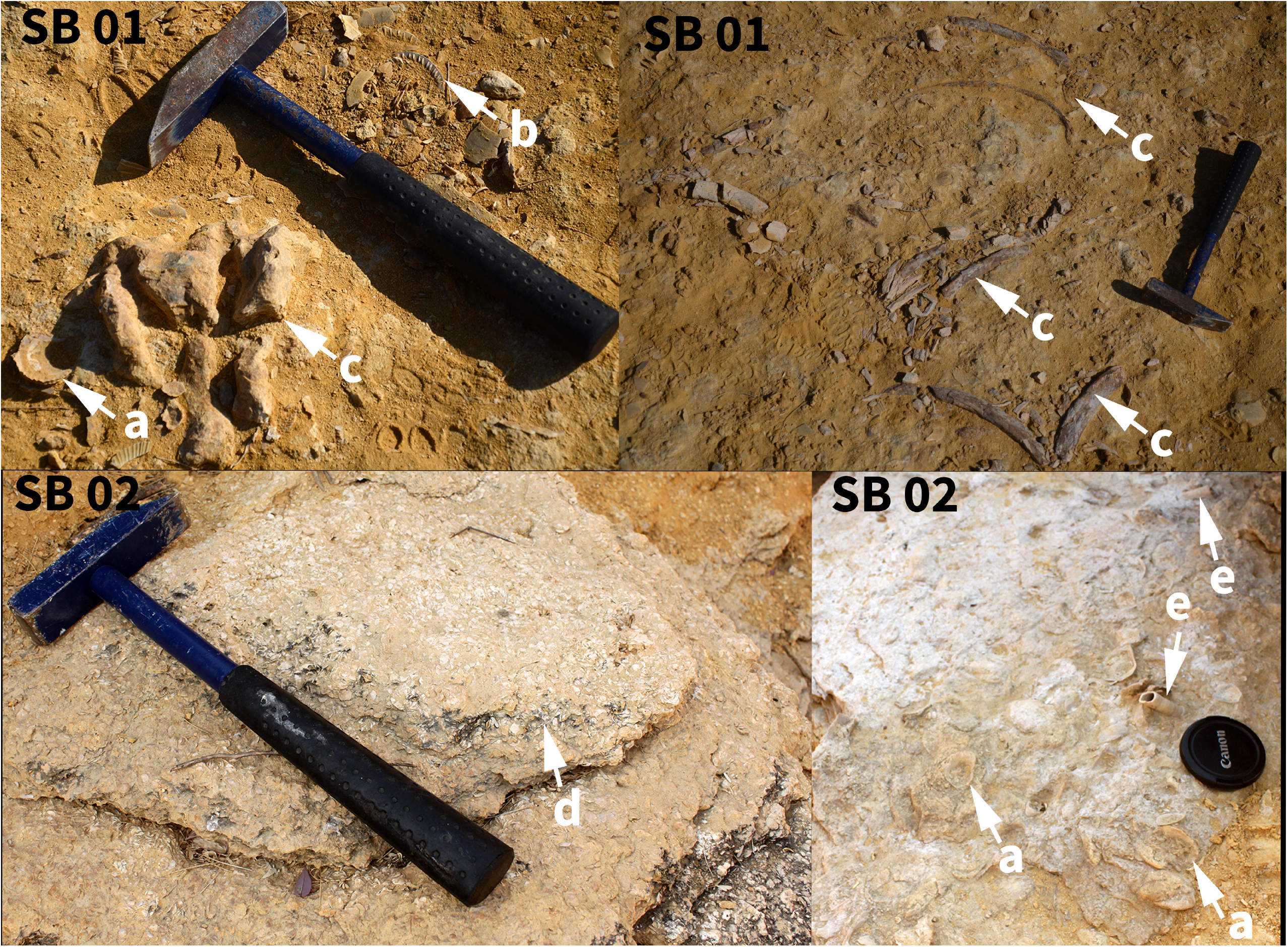
Field photographs depicting diverse fauna of the Oligocene shellbeds of the Kutch basin from Maniyara Fort Formation. It yielded a rich variety of faunal remains such as shells of Ostreidae bivalve (a), Pectinidae bivalve (b), vertebrate bones (c), larger foraminifera (d), Teridinidae bivalve (e).

**Fig 4.**
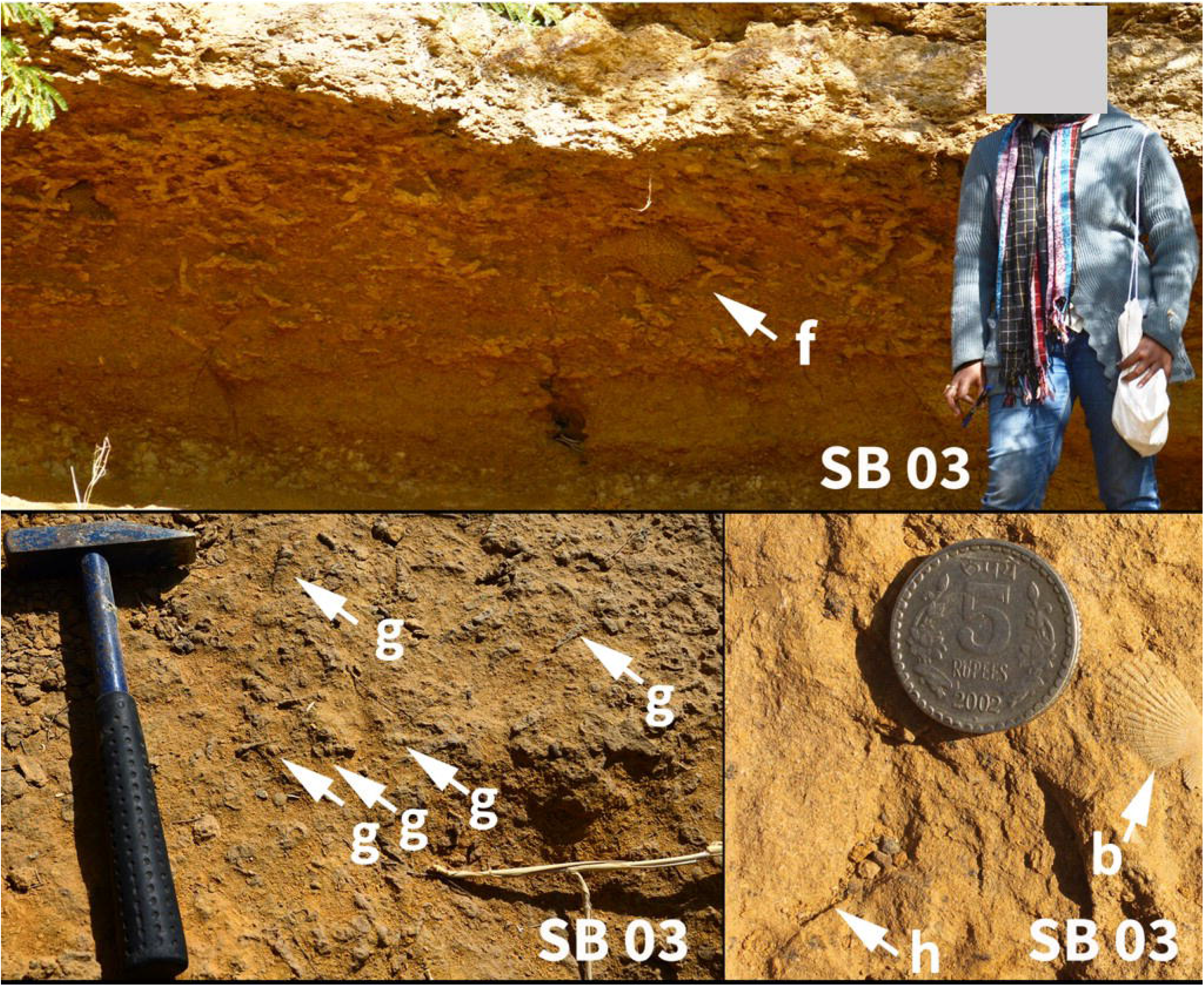
Field photographs depicting diverse fauna of the early Miocene shellbeds of the Kutch basin from Khari Nadi Formation. It yielded a rich faunal composition such as extensive network of burrows (f), Turritellidae gastropod (g), burrows (h) and Pectinidae bivalve (b).

**Fig 5.**
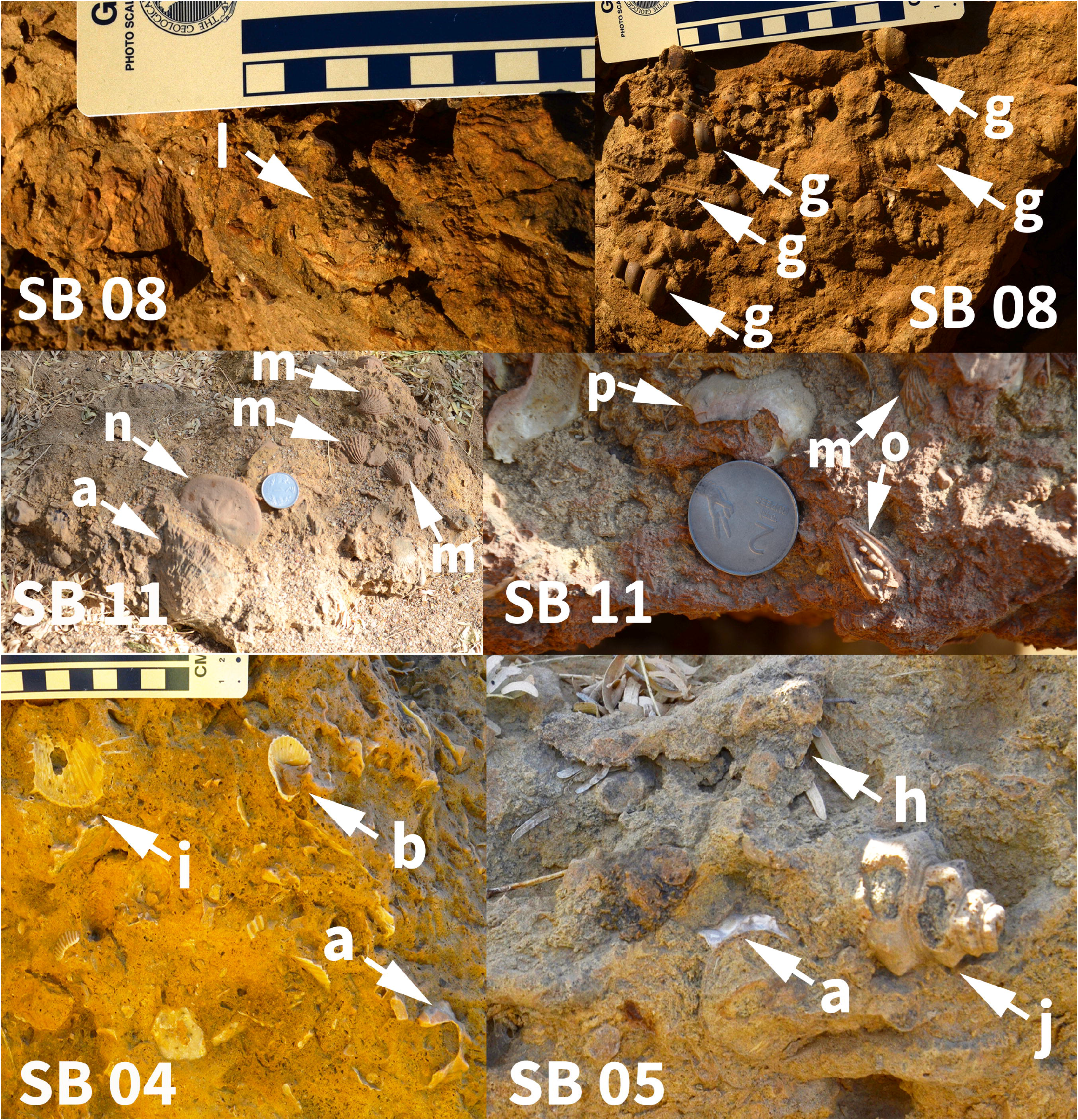
Field photographs depicting diverse fauna of the early Miocene shellbeds of the Kutch basin from Chhasra Formation. Chhasra Formation yielded a rich fauna composed such as Spondylidae bivalve (i), Strombidae gastropod (j), regular echinoid (l), echinoid spine (m), Carditidae bivalve (n), irregular echinoid (o), decapod claw (p), Anomiidae bivalve (q) in addition to (a), (b), (g) and (h).

**FIG 6.**
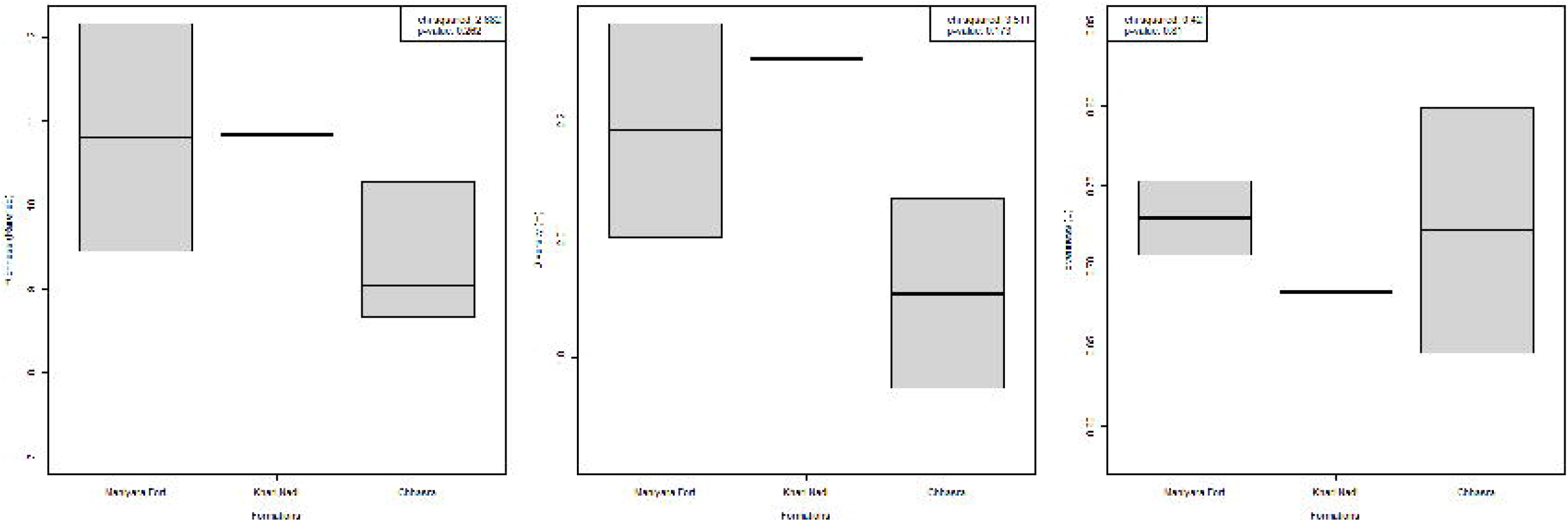
Diversity profile of Oligocene-early Miocene bivalve fauna of Kutch, India. The panel represents comparison of A) rarefied richness, B) Shannon’s diversity index (H) and C) Shannon’s evenness index (J) across formations. The boxes are defined by 25th and 75th quantiles; thick line represents the median value. The individual values are represented by black circles.

Overall, most of the fossils in the shellbeds are calcitic with an exception in SB 02 shellbed. Bivalves are preserved as whole large fragments (umbo included). Aragonite shells are often preserved articulated with some possessing the original shell minerology with well preserved external shell structure (Carditidae), while others are preserved as moulds (e.g. primarily Veneriidae). Calcitic shells, even though mainly disarticulated, are characterized by exceptional preservation of shell ornamentation and dentition. The well-preserved bivalve assemblages suggest minimal transport and physical disruption of fossil material. However, dominance of calcitic forms in many of the shellbeds indicate strong diagenetic alteration potentially influencing the faunal record.

### Collection and identification

We collected bivalve shells from the shell beds by surface sampling during field trips conducted over five years (2011-16). Where the shells were strongly attached to the substrate and posed a threat of breakage during collection, we took detailed photographs for further documentation. Specimens that are represented by 70% of the original shell includingthe umbo were considered for identification and size measurements. Our collection of 2252 individuals represent a total of 15 families and 61 morphospecies (Table 2).

**Table 2.**
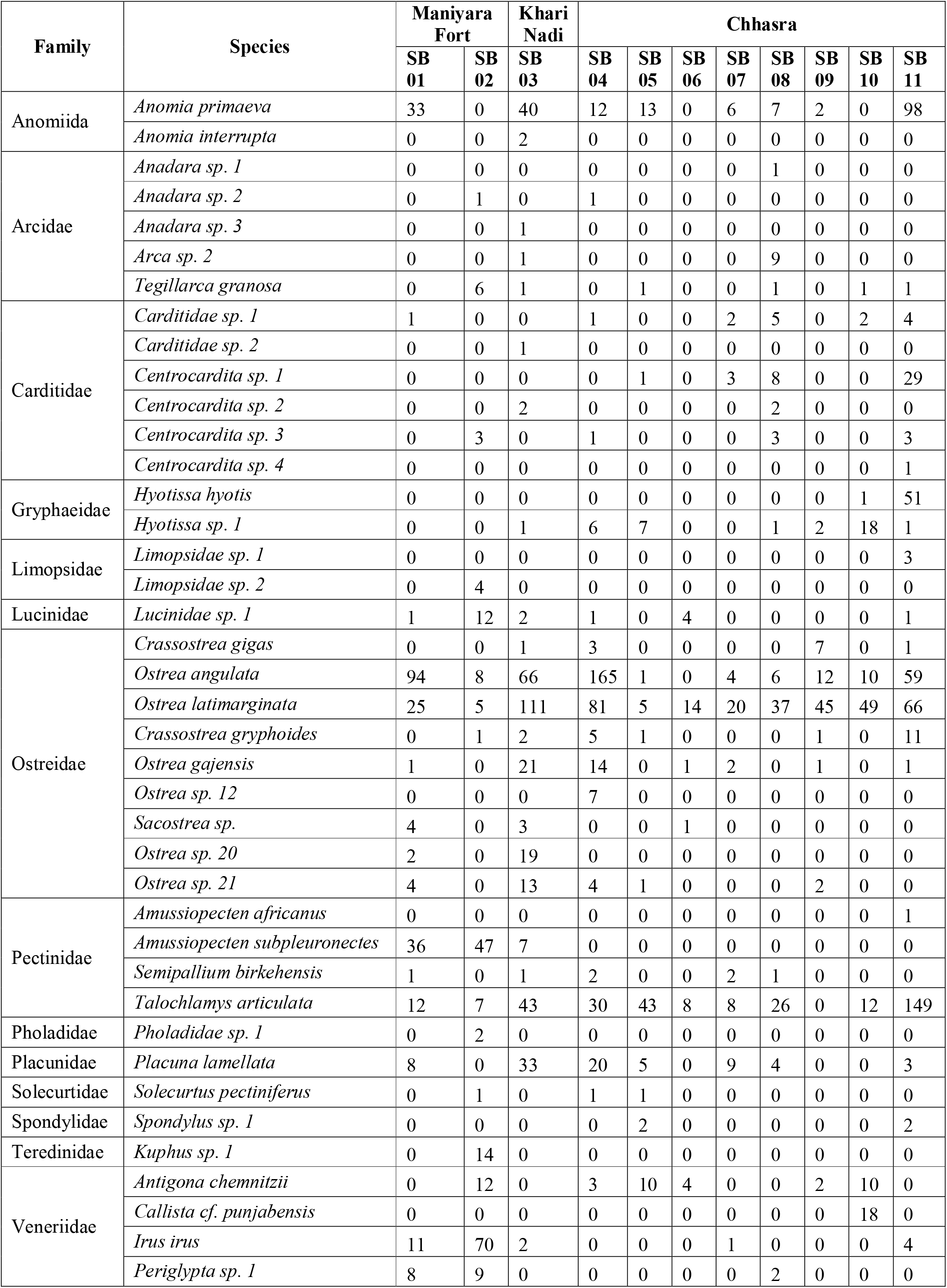
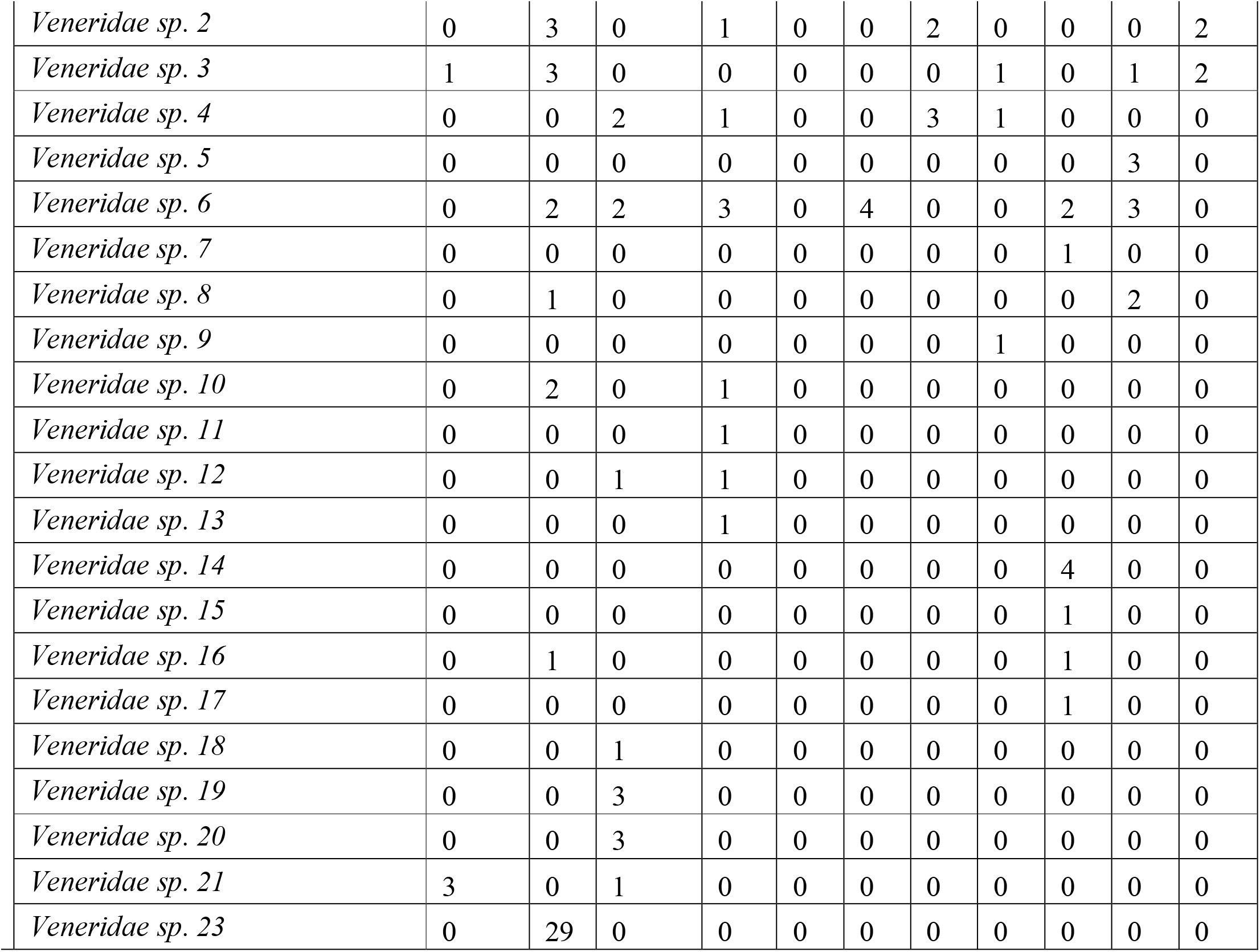
Summary of the bivalve specimens from the shellbeds of Oligocene - early Miocene formations of the Kutch Basin.

The bivalve specimens are identified to lowest possible taxonomic level using the monographs by Vredenburg (1928), Eames and Cox (1956), Sowerby (1840), d’Orbigny (1852), Cox (1927) and other published literature. Although, we could establish the species identity of many of the specimens, we performed our analyses using morphospecies (Campbell and Valentine, 1977) to include all specimens. Information about life habits which included the substrate preference for each bivalve family, was acquired from Pbdb (http://fossilworks.org).

### Measuring body size

We measured the maximum length and height of the specimens using the image processing software (ImageJ 1.47v) to the nearest 0.01 mm. We used log_2_(√(length * height)) as a measure of the body size for each individual following well accepted convention (Kosnik et al., 2006, Chattopadhyay et al, 2015). The body size has been calculated for all the species of each shell bed.

### Analyses

We evaluated the diversity using rarefied species richness (at sample size = 38), Shanon’s diversity index (H) and Shanon’s evenness (J) for each shellbed and specific formations. We used Kruskal-Wallis rank sum test to compare these measures across the three formations. We evaluate the proportional abundances of the calcitic taxa to asses the relative degree of diagenesis affecting preservation.

Two-dimensional ordination assemblages were created with principal coordinate analysis (PCoA), which attempts to represent the pairwise dissimilarity between the lithounit-wise species composition in a low-dimensional space. PCoA was performed on the proportional abundances of the species using capscale functions in the vegan package in R.

We assessed the tempo of the change in the ecological metrics (rarefied richness and overall size) using the maximum likelihood estimation (MLE) model selection fitted to three basic models of temporal change using the paleoTS package: (1) change without directional bias (unbiased random walk – URW), (2) change with directional bias (general random walk – GRW), and (3) no net change (stasis). We have used the reported ages of five shellbeds and inferred ages for the remaining shellbeds based on their stratigraphic position for the MLE analysis. We followed the protocol proposed by Hunt (2006, 2008) and used the paleoTS package in the R programming environment. All statistical analyses were performed in R version 1.1.383 (R Core Team. 2012).

## RESULTS

### Faunal changes

We observed a monotonic, although not significant, increase in the rarefied species richness and a decrease in evenness from the Maniyara Fort (Bermoti Member) to the Chhasra Formation (Kruskal-Wallis rank sum test: H= 0.28, p = 0.87) (Fig. 6, Table 3).

**Table 3.** Ecological metrics measured and estimated for the bivalve assemblages in the Oligocene - early Miocene formations of the Kutch Basin.

| Formation | Unit | Specimen | Raw richness | Shellbed specific |  |  | Formation specific |  |  |
| --- | --- | --- | --- | --- | --- | --- | --- | --- | --- |
|  |  |  |  | Rarefied richness | Shannon diversity index (H) | Evenness (J = H/Hmax) | Rarefied richness | Shannon diversity index (H) | Evenness (J = H/Hmax) |
| Chasra | SB 11 | 493 | 22.0 | 8.5 | 2.0 | 0.65 | 32.71 | 2.43 | 0.63 |
|  | SB 10 | 130 | 13.0 | 9.0 | 1.9 | 0.76 |  |  |  |
|  | SB 09 | 84 | 15.0 | 9.8 | 1.7 | 0.64 |  |  |  |
|  | SB 08 | 116 | 18.0 | 11.3 | 2.2 | 0.75 |  |  |  |
|  | SB 07 | 62 | 12.0 | 10.7 | 2.1 | 0.85 |  |  |  |
|  | SB 06 | 36 | 7.0 | 7.0 | 1.6 | 0.84 |  |  |  |
|  | SB 05 | 91 | 13.0 | 8.8 | 1.8 | 0.69 |  |  |  |
|  | SB 04 | 366 | 25.0 | 9.1 | 1.9 | 0.58 |  |  |  |
| Khari Nadi | SB 03 | 386 | 29.0 | 10.8 | 2.3 | 0.68 | 29 | 2.30 | 0.68 |
| Manyara Fort | SB 02 | 243 | 23.0 | 12.2 | 2.4 | 0.75 | 30.12 | 2.55 | 0.73 |
|  | SB 01 | 245 | 17.0 | 9.5 | 2.0 | 0.71 |  |  |  |

The species association is well distributed across the formations, except for a few unique species. *Kuphus sp. 1* is restricted to the Maniyara Fort Formation. *Amussiopecten* sp. has never been found in the Chhasra Formation. Two species, *Spondylus sp. 1* and *Hyotissa hyotis*, occur exclusively in the Chhasra Formation. The five most abundant species— *Ostrea latimarginata, Ostrea angulata, Talochlamys articulata, Anomia primaeva* and *Placuna lamellata*, are represented in all the formations (Fig. 7). Except for *Ostrea angulata*, all other species show an increase in their relative abundance from the Maniyara Fort to the Chhasra Formation (Table 3). The family Veneridae, in contrast, shows the most dramatic drop in relative abundance (Table 3).

**FIG 7.**
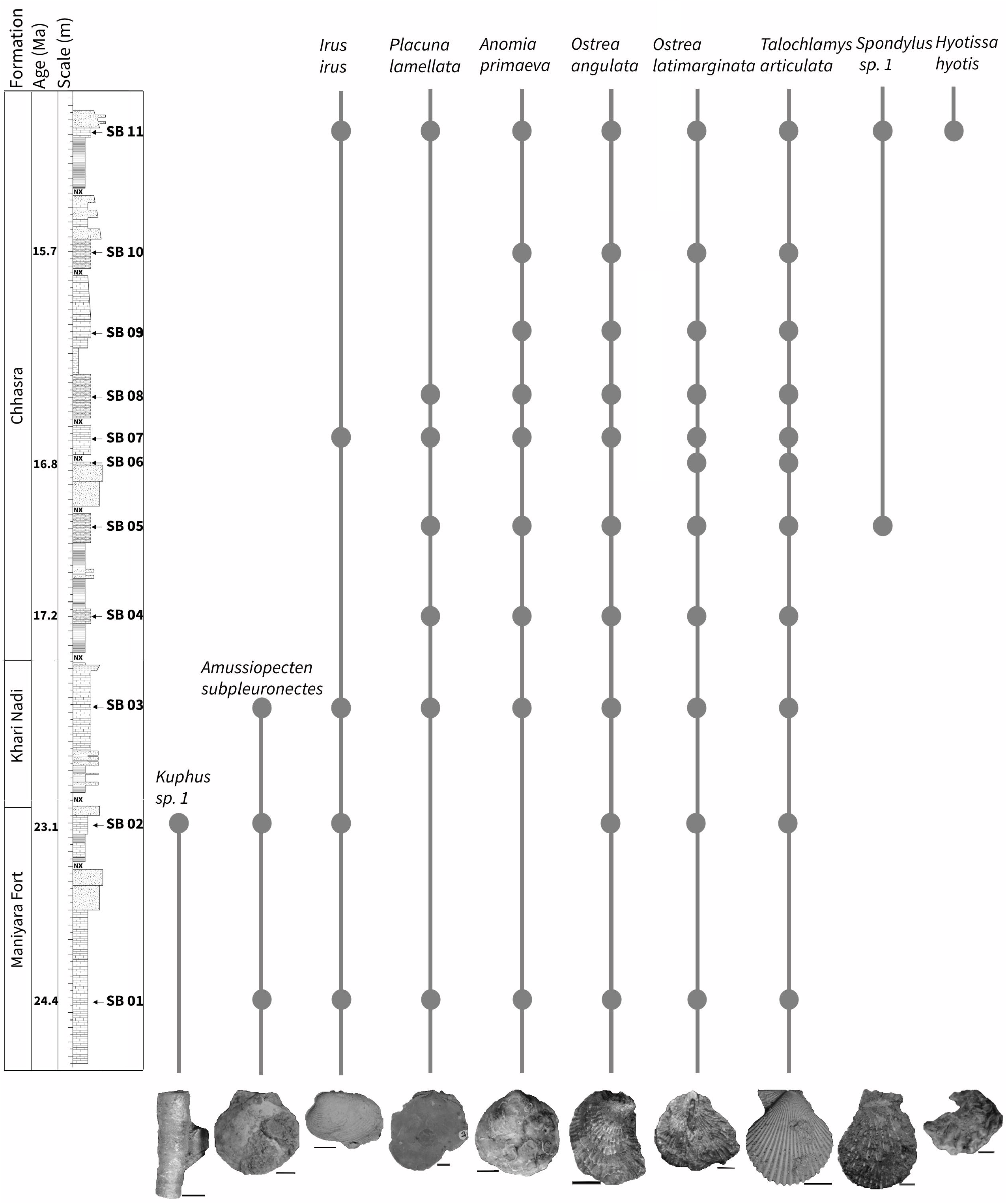
Stratigraphic range of bivalve species in Oligocene - early Miocene succession of the Kutch basin, with the lithostratigraphic, biostratigraphic and Sr-stratigraphic zonations of Biswas et al., 1992 and Dutta et al., 2020. The recorded occurrences at each shellbed units are represented by solid horizontal lines. The photograph at the bottom of the graph represents the image of the bivalve species which are listed in the graph. From left to right: *Kuphus sp*.*1, Amussiopecten subpleuronectes, Irus irus, Placuna lamellata, Anomia primaeval, Ostrea angulata, Ostrea latimarginata, Talochlamys articulata, Spondylus sp*.*1, Hyotissa hyotis*.

The species composition shows two clusters in the PCoA plot, primarily separating Oligocene shellbeds (Maniyara Fort Formation) from Miocene ones (Khari Nadi and Chhasra formations) for analysis including all species (Fig. 8A), only calcitic species (Fig. 8B), and analysis excluding species of veneriids. The relative distance between the shellbeds of Khari Nadi and the Chhasra Formation changes when some groups are excluded (Fig. 8B-C).

**FIG 8.**
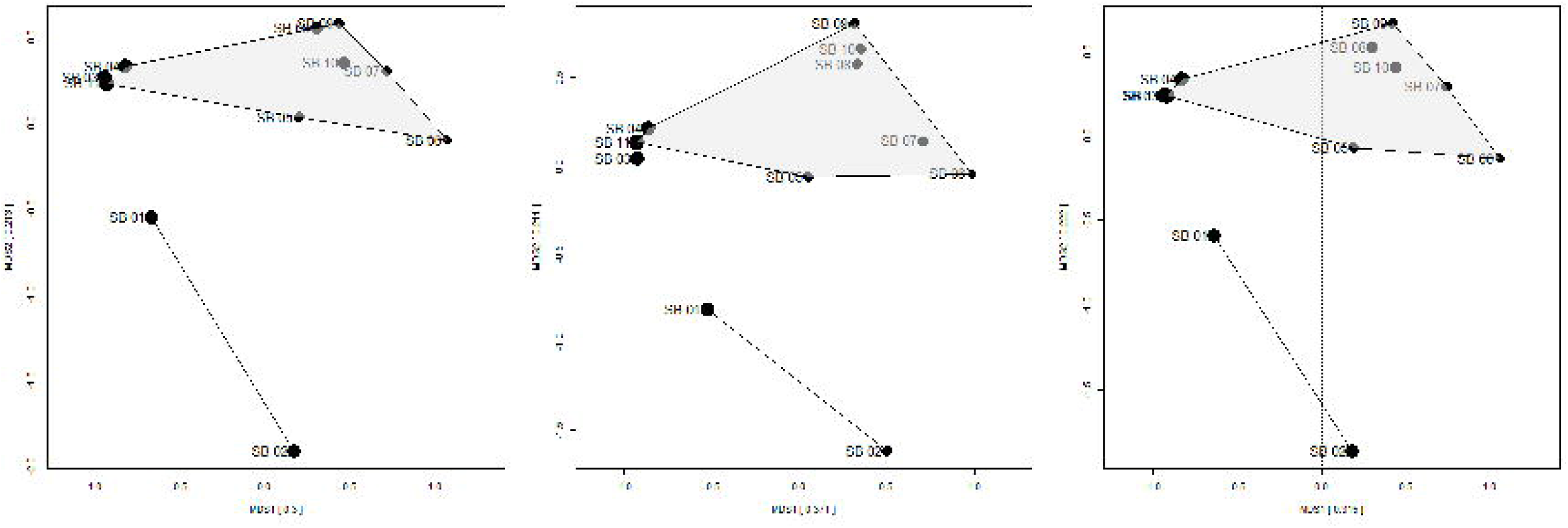
PCoA plot demonstrating the bivalve composition of the shellbeds of Oligocene - early Miocene of the Kutch basin for A) all species, B) all calicitic species, C) species excluding Veneriids.

Although there are changes in the rarefied richness over time (Fig. 9), there is no net change in diversity over time as supported by the MLE analysis. The stasis model (highest Akaike.wt) best explains our data as compared to the other models (Table 4). This result does not change when the analysis is performed by considering only calcitic species (Fig. 9D-F), and excluding species of veneriids (Fig. 9G-I) (Table 4).

**Table 4.** Results of the maximum likelihood estimation (MLE) analyses for detecting temporal change in species-rarefied richness and overall size. The model with the highest weight (Akaike.wt) is most consistent with the data. ΔAICc, difference in AICc from the best-supported model; wAICc, model weights.

|  | **Model** | **AICc** | **$\Delta\text{AICc}$** | **wAICc** |
| --- | --- | --- | --- | --- |
| Rarefied richness<br>(All species) | GRW | 48.0944 | 4.12323 | 0.063 |
|  | URW | 44.1797 | 0.2086 | 0.444 |
|  | Stasis | 43.9711 | 0 | 0.493 |
| Rarefied richness<br>(All calcitic species) | GRW | 45.62 | 5.78 | 0.04 |
|  | URW | 41.70 | 1.86 | 0.27 |
|  | Stasis | 39.84 | 0.00 | 0.69 |
| Rarefied richness<br>(All except species of<br>Veneriidae family) | GRW | 52.83 | 11.13 | 0.00 |
|  | URW | 48.90 | 7.21 | 0.03 |
|  | Stasis | 41.70 | 0.00 | 0.97 |
| Size<br>(All species) | GRW | 50.75 | 6.72 | 0.03 |
|  | URW | 46.83 | 2.80 | 0.19 |
|  | Stasis | 44.03 | 0.00 | 0.78 |

**FIG 9.**
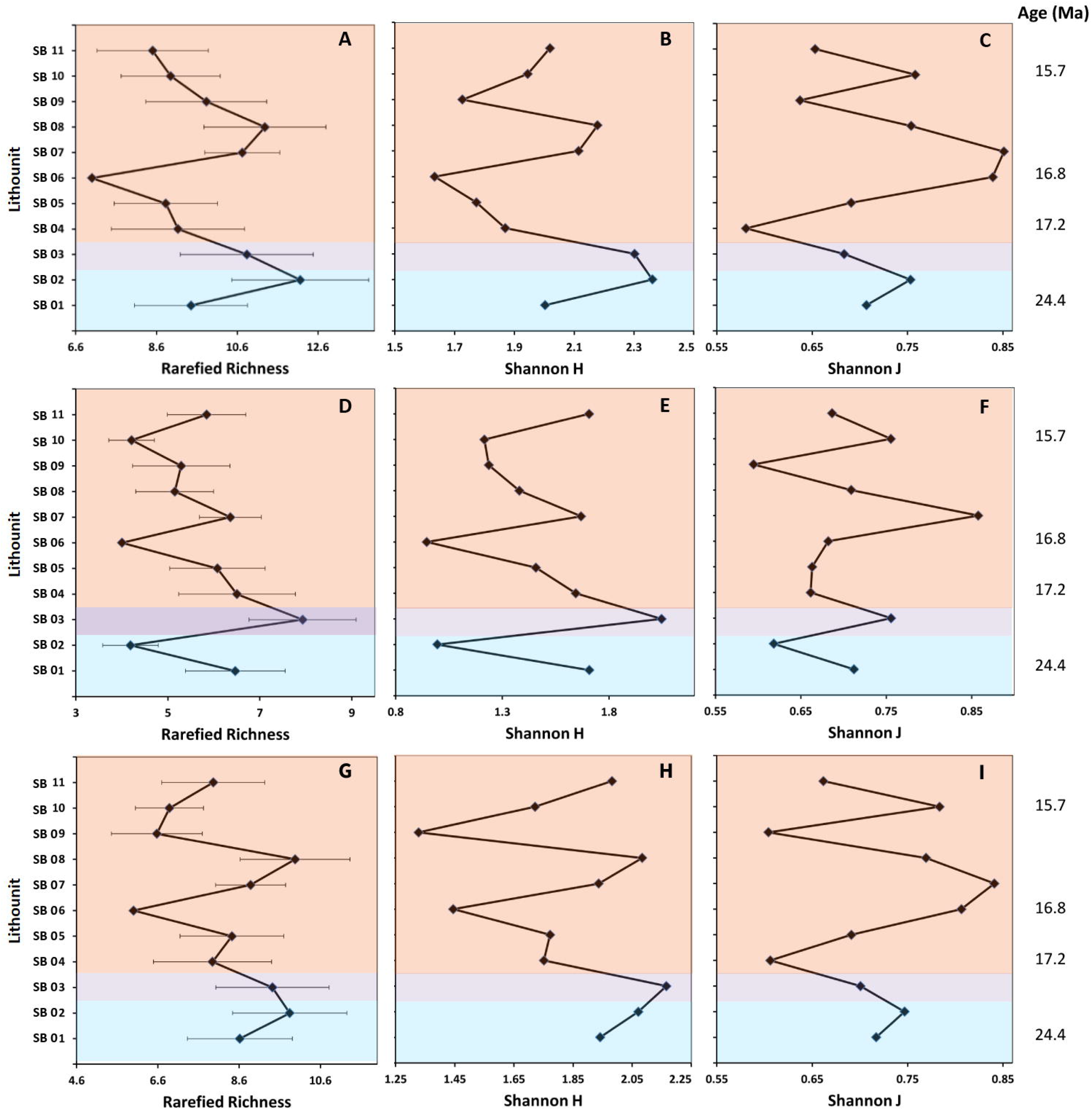
Changes in various measures of community structure over time for all species (A-C), only calcitic species (D-F), excluding Veneriids (G-I). The error bars in the first column (A, D, G) represent the standard deviation (1σ) of rarefied richness for each lithounit. The rightmost label represents the estimated ages for the corresponding shellbed units (Dutta et al., 2020). The background color within each plot represents the formation.

The overall body-size profile through time does not represent any monotonic change across the shellbeds or the formational boundaries (Fig. 10). Eventhough we observe an overall body-size decrease twice – between SB 01-SB 04 (17.2Ma) and SB 05-SB 08 (Fig. 10), the MLE suggests stasis as the best model implying minimal directionality in the net change of the body size for pooled data and species specific analyses (Table 4).

**FIG 10.**
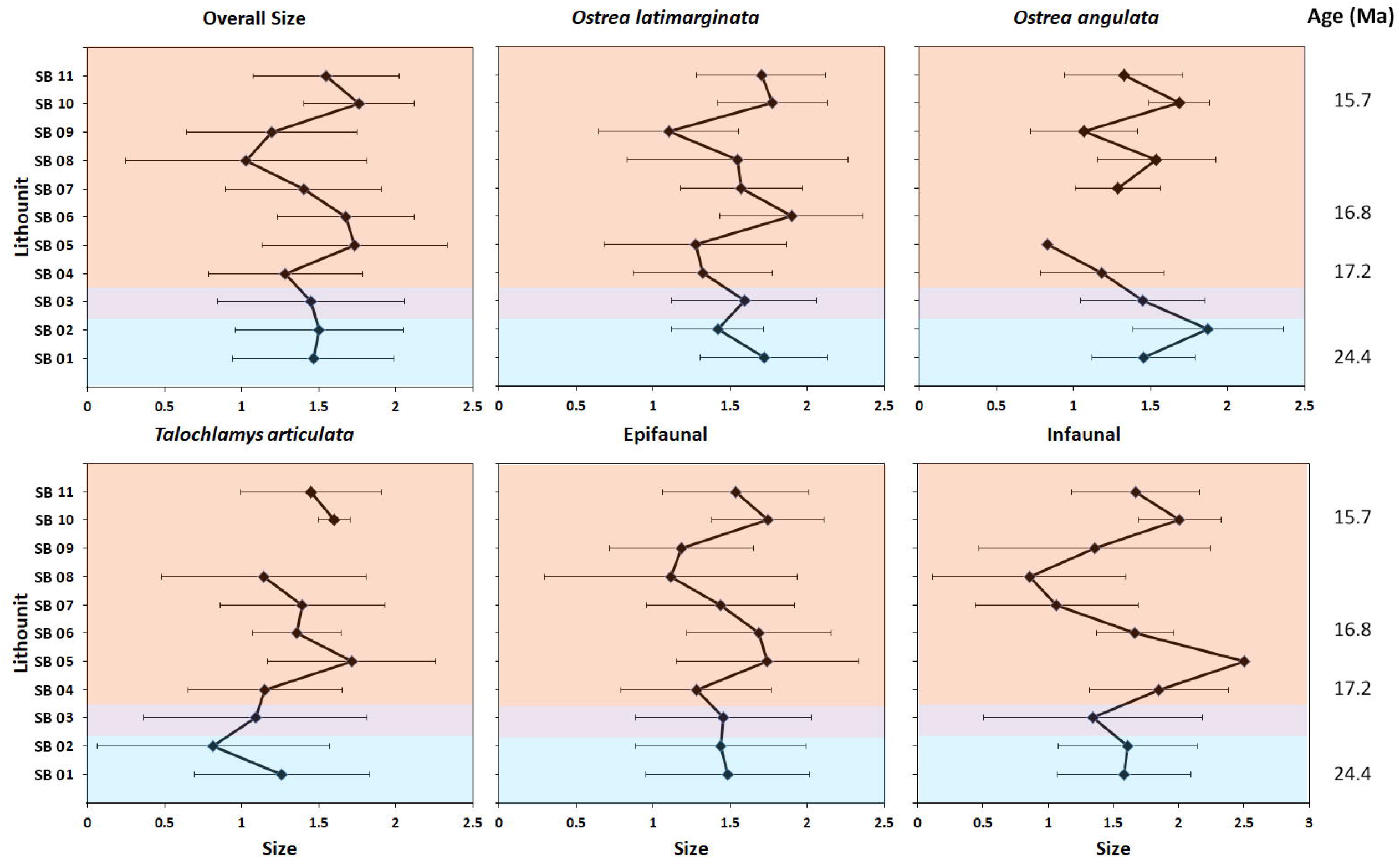
Changes in body size in **A**) overall community, **B**) epifauna, **C**) infauna, **D**) *Talochlamys articulata*, **E**) *Ostrea latimarginata* and **F**) *Ostrea angulata*. The rightmost label represents the estimated ages for the corresponding shellbed units (Dutta et al., 2020). The error bars represent the standard deviation (1σ) of size under each category for each lithounit. The background color within each plot represents the formation.

## DISCUSSION

The changes of the Tethyan seaway during the Oligo-Miocene represent a classic case of tectonic shift that also influenced the regional climate of the Indian subcontinent. This interval witnessed two major regional events, the closure of the Tethys due to development of *Gomphotherium*-Landbridge leading to separation of the Arabian Sea from the proto-Mediterranean Sea (∼20 Ma) and the significant uplift of the Tibetan plateau (∼25 Ma), possibly marking the initiation of monsoon intensification (∼22 Ma) (Bialik et al., 2020, Guo et al., 2008). In a detailed summary of the biotic response to Tethyan seaway closure, Harzhauser et al. (2007) emphasized the practical hurdles of establishing global biogeographic patterns that are often obscured by regional stratigraphic incompleteness, patchy fossil records and preservation biases. They also recognized the need for detailed studies from areas with a relatively poorer record, such as western India. Our study tries to fill this gap by evaluating the nature of the bivalve fauna using well dated shellbeds from the eastern side of the closure and also investigating the role of regional environment in shaping the faunal changes.

### Taphonomic attributes

Before making any inference about faunal changes, it is essential to note the influence of preservation issues, including the taphonomic biases that may have contributed in developing the observed pattern. Here we discuss a few of such issues that potentially affected the faunal record.

In our samples, the role of taphonomic effects in the form of physical damage to the specimens due to transport can be considered minimal. Many of the shells of aragonitic taxa such as Veneridae and Carditidae are preserved articulated and are exquisitely preserved with respect to dentition and external ornamentation (Fig. 11A-B). We also find well-preserved delicate fossils in the majority of the lithounits (Table 1), indicating lack of selective physical removal. Moreover, flat, circular and thin-shelled bivalves such as *Placuna lamellata* are often found oriented parallel to bedding and very well-preserved (Fig. 11C). Quick burial of the shells due to a high rate of sedimentation might have led to this preservation pattern, which further strengthens our claim of in situ preservation with minimal lateral transport.

**FIG 11.**
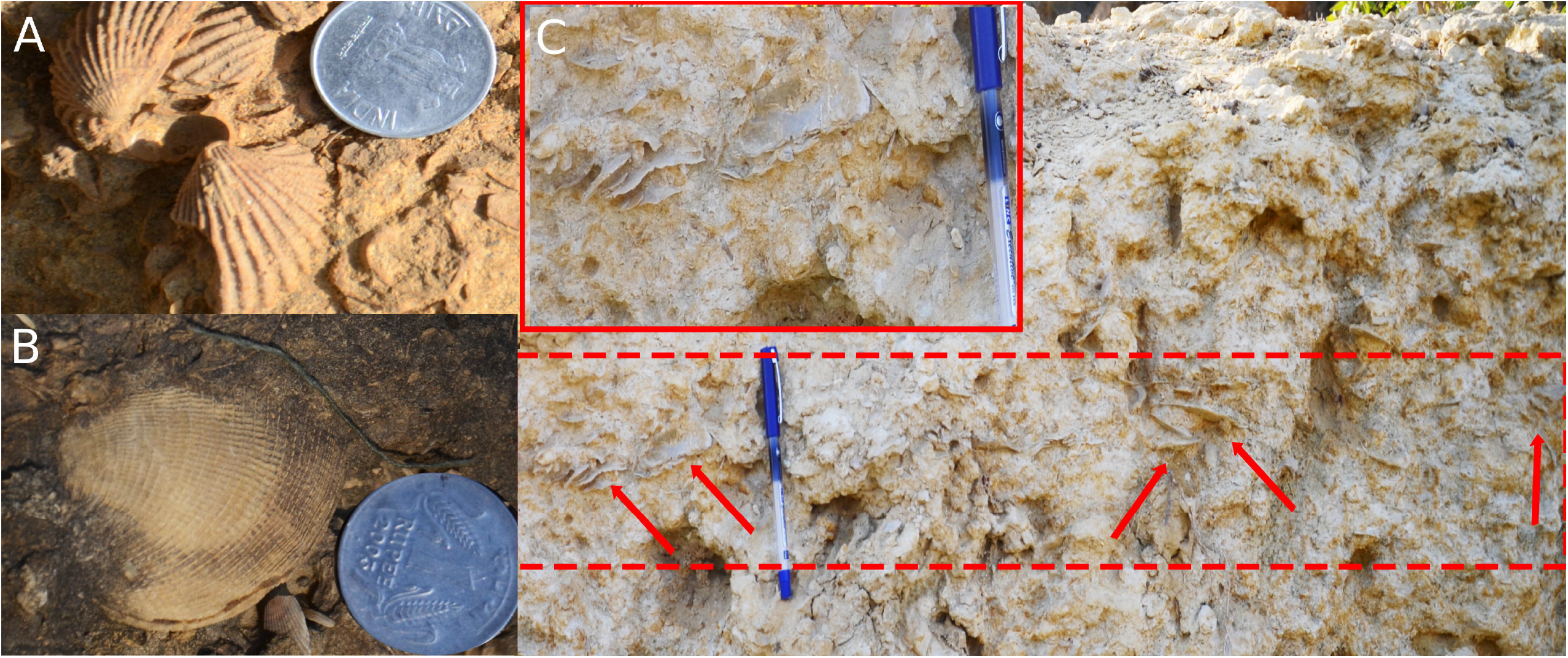
Field photograph indicating the degree of post-mortem transport. Articulated shells of A) an individual of Cardiidae famile and B) an individual of Veneriidae family. These specimens with good preservation indicates limited degree of post-mortem transportation. C) Flat, thin shells of *Placuna lamellata* lying parallel to the beds with excuisite preservation (inset) implying limited transportation. The arrows mark the exact position of some of the shells.

We observed a consistent dominance of calcitic taxa over aragonitic shells (80%) and this trend does not vary significantly across formations (Fig. 12A). An exception to this pattern was observed in SB 02 shellbed. SB 02 shellbed represents a storm event 23.1 My ago which brought infaunal bivalve species (most possessing aragonite shells) from deeper waters to the shallow regime. Overall dominance of calcitic forms (primarily epifaunal), however, suggests limestone deposition during carbonate rich times where the interplay of diagenetic loss of aragonite and the original ecology makes the odds for sampling calcitic taxa eight times higher (Foote et al., 2015). Although such effects would have had an impact on overall diversity, it is less likely to create a trend in the diversity pattern through time. Lack of significant correlation between proportion of calcitic taxa and rarefied richness in various shellbeds (Fig. 12B) supports this. Moreover, species-specific body size is not likely to be affected by selective removal of aragonitic forms. The strong correlation between species-specific body size of the most abundant species (*Ostrea latimarginata*) and body size in other categories (such as total fauna, epifauna, infauna) also corroborates this idea.

**FIG 12.**
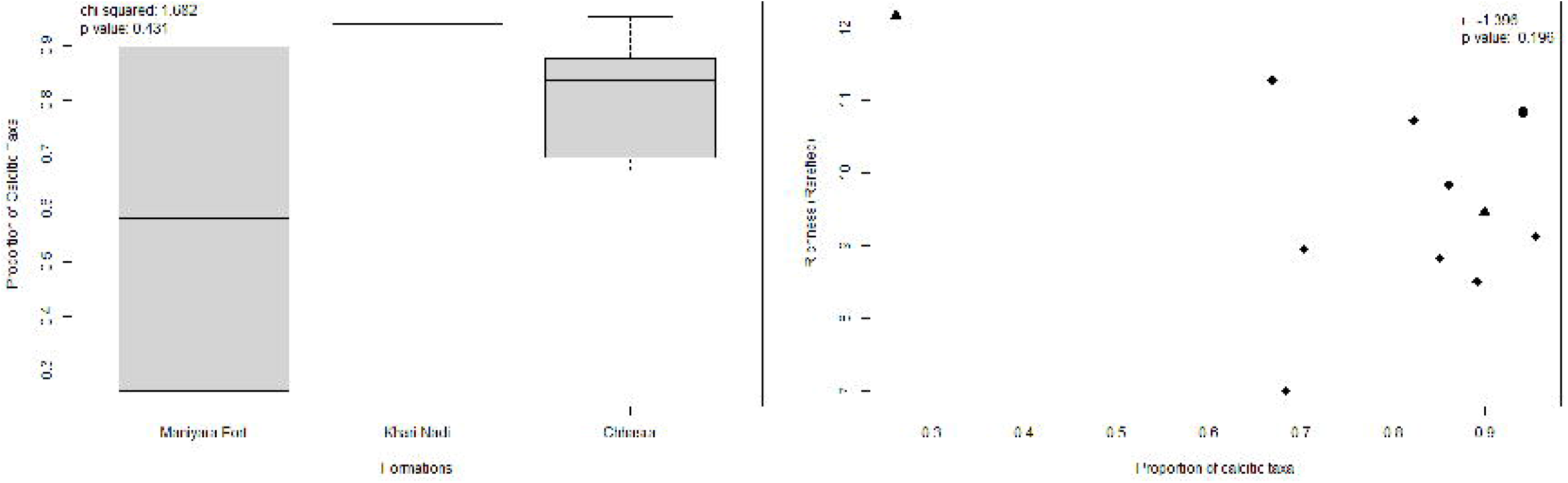
A comparison of proportion of calcitic taxa A) between formations and B) its relationship with the rarefied richness. The boxes are defined by 25th and 75th quantiles; thick line represents the median value. Different geometrical shapes of the points in B represent different formations; triangle, circle and diamond represent Maniyara Fort, Khari Nadi and Chhasra respectively.

### Faunal changes through time

The taxonomy of the Oligo-Miocene molluscan fauna of the Kutch region has been studied by a number of researcher beginning in the 19^th^ century (Sowerby 1847; Vredenberg 1928; Borkar et al. 2004; Borkar et al. 2014; Kulkarni et al. 2010; Halder 2012). Only a few studies explored the paleoecology of this fauna (Kulkarni et al. 2007; Chattopadhyay & Dutta 2013; Bardhan et al. 2014; Patel 2014). The detailed diversity dynamics of this fauna, however, has not been studied before.

The faunal composition of the Kutch Basin changed from Oligocene and maintained a distinct character throughout the early Miocene as revealed by the relatively low variation in species composition through the shellbeds. Although the species association does not vary significantly, there are a few unique associations, such as *Kuphus sp*. in the Maniyara Formation, *Spondylus* sp. 1 and *Hyotissa hyotis* in the Chhasra Formation. Similarly, *Amussiopecten* sp. never appeared in the Chhasra Formation. Some families — Limopsidae, Pholadidae, Tellinidae and Teredinidae — are not present after the Maniyara Fort Formation. The five most abundant species— *Ostrea latimarginata, Ostrea angulata, Talochlamys articulata, Anomia primaeva* and *Placuna lamellata* are present in all the formations implying a steady faunal composition of the shellbeds across the formations (Fig. 7). A shift in the faunal composition from Oligocene shell beds from the younger shellbeds (Fig. 8) is consistent with a seaway closure at the end of Oligocene. The diversity (taxonomic and body size), however, does not show any major shifts throughout the interval. The significant lack of a strong monotonic change supports a largely conservative nature of diversity variation with little directional change as revealed by the MLE analyses (Fig. 9A).

### Paleobiogeographic implication

Previous studies on the biogeography of the vertebrate fauna of Kutch Basin, including fishes (Tewari 1959), reptiles (Head et al. 2007) and marine mammals (Bajpai and Domning 1997; Bajpai et al. 2006; Thewissen and Bajpai 2008, 2009) and terrestrial mammals (Sahni and Mishra 1975; Mishra 1976; Bhandari et al. 2010; Patnaik et al. 2014) demonstrated faunal similarity with north Africa and Europe. Hence, they support events of faunal exchange caused by the opening of the land route between the Afro-Arabian plate and Eurasia during early Miocene. However, Sahni and Mishra (1975) also found evidence of marine incursions that intermittently disrupted this land connection as demonstrated by abundant shark teeth.

Our study shows contradictory patterns of the compositional faunal shift. The documented fauna demonstrates a separation of the cluster composed of the shellbed units of the Maniyara Formation and the rest of the Miocene shellbeds, which signifies a change in species composition across the Oligocene-Miocene boundary (Fig. 8). Taphonomy alone can not explain this results because the results do not change when the analyses is restricted to only calcitic species and non-veneriid species. The timing of this change can be constrained between 23.1Ma (SB 02) and 17.2Ma (SB 04). The proposed age of the Tethyan closure (∼19Ma) falls within this time window and the observed change in species composition is consistent with the expected drop in faunal exchange due to the seaway closure. However, we do not observe distinct cluster in the composition between the bivalve families of WIP, African-Arabian and proto Mediterranean from the Chattian to Burdigalian. The bivalve composition of Chattian shellbeds (SB 01 and SB 02), even though different from the early Miocene shellbeds of the Kutch Basin, demonstrates a significant absence of typical MIP bivalve fauna such as *Loripes, Corbula* and *Cyclocardia* (McCall et al. 1994; Bernasconi and Robba 1993; Mandic and Steininger 2003; Reuter et al. 2012; Zunino and Pavia 2009; Cox 1936; Cherchi et al. 2000). This implies a drop in faunal exchange as early as ∼24.4Ma (Chattian). Such absence of characteristic MIP faunal elements might have been influenced by the lack of aragonitic forms in our assemblage and limits the validity of a global comparison. However, presence of many of these forms from older formations of similar taphonomic grade from the basin (Vredenberg, 1928) points towards the true absence of these forms in the Oligocene formations.

A comprehensive study on the biogeography of gastropods from this area was conducted by Harzhauser et al. (2009), in which he showed endemism starting as early as the Aquitanian. Using a global biogeographic survey of marine gastropods, Harzhauser et al. (2007) demonstrated that the WIP faunas had 27% taxa common with the MIP faunas in the north-eastern Tethyan coast during the Oligocene. This pre-landbridge pattern for gastropods changed completely during the early Miocene, showing a drop in faunistic similarities by 5% between the Pakistan faunas and those from the Proto-Mediterranean Sea (Harzhauser et al. 2002, 2007). According to them, such dramatic change in the WIP probably took place in the Aquitanian and predates the established age of the closure of the Tethyan Seaway (∼19Ma). The gastropod data recorded from the Gaj Formation of Pakistan (Vredenburg, 1925–1928) also show an increasing similarity with the SE Asian fauna and a diminishing resemblance of WTR characters in the early Miocene. However, the lack of precise ages of the gastropod bearing horizons from western India in these studies (Harzhauser et al. 2009) made it difficult to narrow down the age of this event. The observed lack of a progressive change within the formations studied by us speaks for a low-diversity marine community that remained unchanged through the early Miocene (post-Chattian) and supports the previous claim of a limited faunal exchange during the early Miocene (Harzhauser et al., 2007). However, the claim of the existence of a seaway connection during the Oligocene based on the high degree of similarity (27%) between WIP and MIP (Harzhauser et al. 2007) might not be entirely correct. Our results suggest a drop in faunal exchange in the late Oligocene (∼24.4Ma), characterized by the absence of typical MIP forms.

The paleobiogeographic record of pectinids, one of the most abundant fossils in late Cenozoic near-shore deposits, provides information about the faunal exchange to the east of the landbridge. The Neogene pectinid record reflects a dramatic renewal of diversity after a prolonged low-diversity spell during the Paleogene (Raines and Poppe 2006) and resulted in distinct early Miocene pectinid assemblages separated by the *Gomphotherium* Landbridge. The only taxon found on both sides of this landbridge, *Crassadoma multistriata* (Poli 1795), was present even before the closure of the Tethys seaway (Eames and Cox 1956) and continued to exist there. We did not find this genus at our locality, even though other pectinids such as *Talochlamys* dominated the Oligocene-Miocene deposits of the Kutch Basin. One of the pectinids of our study, *Amussiopecten* has been reported from Iran (Qom Formation, Reuter et al. 2009), Pakistan (Gaj Formation, Vredenburg 1925, Iqbal, 1980) and Egypt (Mandic and Piller 2001). We did not encounter *Amussiopecten* in the Kutch deposits after the Aquitanean. The genus *Hyotissa* appeared exclusively in the Chhasra Formation and occurred in the early Burdigalian succession of Egypt (Mandic and Piller 2001). The occurrence of common genera in coeval formations from the western Indian and Arabian-African region, points toward a faunal exchange within the eastern Tethyan seaway in the early Miocene.

Bivalve-based paleobiogeographic reconstruction, such as the one attempted by the present study, although important, might not be sufficient to understand the extent of the seaway connection and regional faunal exchange. It would, therefore, require multi-taxon analyses from time constrained lithounits of this area to fully resolve the extent of faunal exchange during Oligocene-early Miocene time.

### Paleoenvironmental implication

Evolution of the Himalays has been attributed to cause major changes in the climatic configuration and consequently the distribution of terrestrial faunas of the Indian subcontinent. Its effect on shallow marine benthos of the Tethyan sea has yet to be investigated in detail. The exact timing of the initiation of monsoon intensification is heavily debated and the sedimentary record of northwestern India, a severely understudied region, may provide the crucial piece of information as predicted before (Tada et al, 2016). The Oligocene-early Miocene time interval is of particular importance for the Asian climate evolution because of the change in atmospheric circulation that evolved into a dominant monsoon system (Sun and Wang, 2005; Guo et al., 2008). The continental monsoon records, however, often fail to indicate the timing of this crucial change because of insufficient age-constraint of the terrestrial record. It has been speculated that the Asian monsoon started as early as Oligocene based on the pollen data from China (Sun and Wang, 2005) and cyclonic storm beds from the Kutch Basin (Reuter et al 2013). The development of aeolian deposits in the Chinese Loes plateaus also indicates an early initiation of the strengthening of Asian Monsoon around or before 22Ma (Guo et al, 2008). The direct records, however, support a much later strengthening of monsoon (Tada et al, 2016). Based on the sediment thickness in the Arabian Sea, Clift et al (2008b) claimed a monsoonal strengthening event between 16 and 10 Ma linking the high erosion rate with high elevation of the Tibetan Plateau. Others, however, support a much later event of monsoon intensification around 10 My ago (Kroon et al, 1991). A biomarker-based study by Zhuang et al (2017) documents an ocean cooling at 11–10 Ma and supports the establishment of monsoonal upwelling in the western Arabian Sea during that time.

The initiation of the monsoon is predicted to result in higher erosion rates resulting in an increase in terrestrial input and probably a decrease in salinity in the shallow coastal region. The resulting changes in salinity and sedimentation rate are expected to affect shallow marine communities. Monsoon-influenced changes in environmental parameters are observed to influence the species composition and diversity of Recent bivalves in the coastal waters of India (Sarkar et al, 2019; Chattopadhyay et al, 2021). It is not unusual to have relatively low diversity in areas subject to stress conditions due to reduced salinity, and a negative correlation is expected between richness and terrigenous input (Careddu et al, 2015). The diversity of the bivalve assemblages during the observed interval, however, has not changed significantly (Fig. 6). Even the species composition remains unchanged through the Budigalian-Langhian interval (Fig. 8) represented by SB 04 (∼17.2 Ma) to SB 10 (∼15.7Ma) – an interval that has been claimed to have experienced the initiation of monsoon (Clift et al, 2008 b).

The major variation in community composition across the Oligo-Miocene also coincides with a facies change. Reuter et al (2013) noted a significant change in sedimentation characterized by an increase in the relative abundance of siliciclastics from the late Oligocene to the early Miocene of Kutch. Our field observation also supports this, with the maximum change occurring at the transition of SB 02 (Maniyara Fort, ∼23.1My) and SB 03 (Khari Nadi). SB 03 is the shellbed with highest proportion of siliciclastic grains among the studied shellbeds and is characterized by clastic sedimentary features such as bedding and trough cross-stratification (Biswas, 1992). Hence, the compositional change across the Oligocene and Miocene (Fig. 8) could even reflect a facies change. The variation in facies composition within Khari Nadi and Chhasra shellbeds, however, is not reflected in the coordination analyses. SB 03 (Khari Nadi) and SB 04 (Chhasra) consistently show higher similarity in species composition inspite of their difference in facies: in contrast to the highly siliciclastic-rich SB 03, the composition of SB 04 is primarily limestone. The facies composition and associated preservational artifact, hence, makes only a limited contribution to explaining the compositional variation of the fauna in the shellbeds; the variation probably reflects the true ecology of the fauna during this interval.

Absence of any temporal trend in body-size in our study supports the existence of a morphological stasis in the community for ∼9Ma. Although detection of the climatic effect on body size might be tricky in relatively short ecological duration (Smith et al., 1995, Berke et al., 2013, Ohlberger et al., 2012, Jablonski, 1997), it is important to note that such an unresponsive nature of shallow marine faunas is not unexpected from a tropical biota. Tropical species are known to show a lower physiological response and hence display a strong niche conservatism (Romdal et al, 2013; Khaliq et al, 2015). Our results are consistent with this claim and point to its existence in the Oligo-Miocene time interval.

## CONCLUSION

Bivalve assemblages, collected through detailed stratigraphic sampling of the Maniyara Fort (Chattian), Khari Nadi (Aquitanian) and Chhasra formations (Burdigalian) provide insight into the nature of marine biota of western India during a crucial time marked by the Tethyan seaway closure and the Himalayan uplift. The bivalves demonstrate a temporal change in species composition across the Oligocene – Miocene boundary. The observed pattern of change in faunal character between 23.1 Ma and 17.2Ma is consistent with the closure of the Tethyan seaway (∼19Ma) and a resultant halt in faunal exchange. The absence of key proto-Mediterranean taxa in Oligocene shellbeds, however, supports a limited faunal exchange even in the Chattian (∼24.4Ma). Absence of any change in either diversity (taxonomic and body size) or species composition during an interval between ∼17.2 Ma to ∼15.7Ma, an interval that has been claimed to have experienced the initiation of monsoon, points to the limited influence of this phenomenon on the shallow marine ecosystem. Although we can not completely rule out the influence of taphonomy on the observed faunal pattern, our results demonstrate little or no influence of the Tethyan closure and Himalayan uplift on the Oligo-Miocene bivalve fauna of the Kutch Basin.

## ACKNOWLEDGEMENTS

We are grateful to Debarati Chattopadhyay, Shibajyoti Das, Rohini Das and Neeti Mandal for their help in data acquisition in the field and in preliminary identification. We would like to thank Oleg Mandic for his help in bivalve identification. We thank Linda Ivany, Martin Zuschin for their comments on an earlier draft. SD is supported by the INSPIRE fellowship. DC acknowledges the financial support of DST-FAST (SR/FTP/ES-85/2012 dated 27.02.2012).

## REFERENCES

Bajpai, S. and Domning, D.P., 1997, A New Dugongine Sirenian from the Early Miocene of India: Journal of Vertebrate Paleontology, v. 17, p. 219–228.

Bajpai, S., Thewissen, J.G.M., Kapur, V.V.I.R., Tiwari, B.N. and Sahni, A., 2006, Eocene and Oligocene Sirenians (Mammalia) from Kachchh, India: Journal of Vertebrate Paleontology, v. 26, p. 400–410.

Bakun, A., Field, D. B., Redondo-Rodriguez, A. N. A., and Weeks, S. J., 2010. Greenhouse gas, upwelling-favorable winds, and the future of coastal ocean upwelling ecosystems: Global Change Biology, v. 16(4), p. 1213–1228.

Bardhan, S., Mallick, S. and Das, S.S., 2014, Palaeobiogeographic constraints on drilling gastropod predation: a case study from the Miocene Khari Nadi Formation in Kutch, Gujarat: Special Publication of the Palaeontological Society of India, v. 5, p. 205–213.

Berke, S. K., Jablonski, D., Krug, A. Z., Roy, K., and Tomasovych, A., 2013, Beyond Bergmann’s rule: size–latitude relationships in marine Bivalvia world-wide: Global Ecology and Biogeography, v. 22, p. 173–183.

Bernasconi, M.P. and Robba, E., 1993, Molluscan palaeoecology and sedimentological features: an integrated appraoch from the Miocene Meduna section, northern Italy: Palaeogeography, Palaeoclimatology, Palaeoecology, v. 100, p. 267–290.

Bhandari, A., Dun, D., Mohabey, D.M., Bajpai, S., Tiwari, B.N., Dun, D. and Pickford, M., 2010, Early Miocene mammals from central Kutch (Gujarat), Western India: Implications for geochronology, biogeography, eustacy and intercontinental dispersals: Neues Jahrbuch für Geologie und Paläontologie, v. 256, p. 69–97.

Biswas, S.K., 1992, Tertiary stratigraphy of Kutch: Journal of the Palaeontological Society of India, v. 37, p. 1–29.

Borkar, V.D., Kulkarni, K.G. and Bhattacharjee, S., 2004, Molluscan fauna from the Miocene sediments of Kachchh, Gujarat, India - Part 1. Oysters: Geophytology, v. 34, p. 1–7.

Borkar, V.D., Kulkarni, K.G. and Bhattacharjee-Kapoor, S., 2014, Molluscan fauna from the Miocene Sediments of Kachchh, Gujarat, India - Part 4, Indarca, a new anadoroid genus: Journal of the Geological Society of India, v. 83, p. 290–294.

Campbell, C.A. and Valentine, J.W., 1977, Comparability of modern and ancient marine faunal provinces: Paleobiology, v. 3, p. 49–57.

Careddu, G., Costantini, M. L., Calizza, E., Carlino, P., Bentivoglio, F., Orlandi, L., and Rossi, L., 2015, Effects of terrestrial input on macrobenthic food webs of coastal sea are detected by stable isotope analysis in Gaeta Gulf: Estuarine, Coastal and Shelf Science, v. 154, p. 158–168.

Catuneanu, O. and Dave, A., 2017, Cenozoic sequence stratigraphy of the Kachchh Basin, India: Marine and Petroleum Geology, v. 86, p. 1106–1132.

Chattopadhyay, D. and Dutta, S., 2013, Prey selection by drilling predators: A case study from Miocene of Kutch, India: Palaeogeography, Palaeoclimatology, Palaeoecology, v. 374, p. 187–196.

Chattopadhyay, D., Zuschin, M., and TomašovýCh, A., 2015, How effective are ecological traits against drilling predation? Insights from recent bivalve assemblages of the northern Red Sea. Palaeogeography, Palaeoclimatology, Palaeoecology, v. 440, p. 659–670.

Chattopadhyay, D., Sarkar, D., Bhattacharjee, M., 2021, The distribution pattern of marine bivalve death assemblage from the western margin of Bay of Bengal and its oceanographic determinants. Frontiers in Marine Science. v. 8, p. 613

Cherchi, A., Murru, M. and Simone, L., 2000, Miocene Carbonate Factories in the Syn-rift Sardinia Graben Subbasins (Italy): Facies, v. 43, p. 223–240.

Clift, P. D., and Webb, A. A. G. (2019). A history of the Asian monsoon and its interactions with solid Earth tectonics in Cenozoic South Asia. Geological Society, London, Special Publications, 483(1), 631–652.

Clift, P. D., Hodges, K. I. P. V, Heslop, D., Hannigan, R., Long, H. V. A. N., and Calves, G. (2008a). Holocene erosion of the Lesser Himalaya triggered by intensified summer monsoon: Geology, v. 36, p. 79–82.

Clift, P. D., Hodges, K. I. P. V, Heslop, D., Hannigan, R., Long, H. V. A. N., & Calves, G. (2008b). Correlation of Himalayan exhumation rates and Asian monsoon intensity: Nature Geoscience, v. 17, p. 875–880.

Clift, P., and Gaedicke, C., 2002, Accelerated mass flux to the Arabian Sea during the middle to late Miocene: Geology, v. 30(3), p. 207–210.

Cox, L.R., 1927, Neogene and Quarternary Mollusca from the Zanzibar Protectorate: Report on the Palaeontology of the Zanzibar Protectorate, p. 13–102.

Cox, L.R., 1936, Fossil Mollusca from southern Persia (Iran) and Bahrein Island: Memoirs of the Geological Survey of India, v. 22, p. 1–67.

D’Orbigny, A., 1852, Paléontologie Francaise: Description des Mollusques et Rayonnés Fossiles de France: Terrains Crétacés, Bryozoaires. Librairie Victor Masson: Paris, v. 5, p. 185–472.

Diniz-Filho, J. A. F., and Bini, L. M., 2008, Macroecology, global change and the shadow of forgotten ancestors: Global Ecology and Biogeography, v. 17(1), p. 11–17.

Dutta, S., Chattopadhyay, D., Chattopadhyay, D., Misra, S., and Turchyn, A. V., 2020, Strontium stratigraphy of the Oligocene–Early Miocene shellbeds of the Kutch Basin, western India, and its implications: Lethaia.

Eames, F.E. and Cox, L.R., 1956, Some Tertiary pectinacea from East Africa, Persia and the mediterranean region: Journal of Molluscan Studies, v. 32, p. 1–6.

Fields, P. A., Graham, J. B., Rosenblatt, R. H., and Somero, G. N., 1993, Effects of expected global climate change on marine faunas: Trends in Ecology & Evolution, v. 8(10), p. 361–367.

Gilbert, D., Rabalais, N. N., Diaz, R. J., and Zhang, J., 2010. Evidence for greater oxygen decline rates in the coastal ocean than in the open ocean: Biogeosciences, v. 7(7), p. 2283.

Guo, Z. T., Ruddiman, W. F., Hao, Q. Z., Wu, H. B., Qiao, Y. S., Zhu, R. X., and Liu, T. S., 2002, Onset of Asian desertification by 22 Myr ago inferred from loess deposits in China: Nature, v. 416(6877), p. 159.

Halder, K., 2012, Cenozoic fossil nautiloids (Cephalopoda) from Kutch, western India: Palaeoworld, v. 21, p. 116–130.

Harzhauser, M., 2007, Oligocene and Aquitanian gastropod faunas from the Sultanate of Oman and their biogeographic implications for the early western Indo-Pacific: Palaeontographica, v. 280, p. 75–121.

Harzhauser, M., Kroh, A., Mandic, O. and Piller, W.E., 2007, Biogeographic responses to geodynamics: A key study all around the Oligo – Miocene Tethyan Seaway: Zoologischer Anzeiger, v. 246, p. 241–256.

Harzhauser, M., Piller, W.E. and Steininger, F.F., 2002, Circum-Mediterranean Oligo–Miocene biogeographic evolution–the gastropods’ point of view: Palaeogeography, Palaeoclimatology, Palaeoecology, v. 183, p. 103–133.

Harzhauser, M. and Piller, W.E., 2007, Benchmark data of a changing sea— palaeogeography, palaeobiogeography and events in the Central Paratethys during the Miocene: Palaeogeography, Palaeoclimatology, Palaeoecology, v. 253, p. 8–31.

Harzhauser, M., Reuter, M., Piller, W.E., Berning, B., Kroh, A. and Mandic, O., 2009, Oligocene and Early Miocene gastropods from Kutch (NW India) document an early biogeographic switch from Western Tethys to Indo-Pacific: Paläontologische Zeitschrift, v. 83, p. 333–372.

Head, J.J., Mohabey, D.M. and Wilson, J.A., 2007, Acrochordus hornstedt (Serpentes, Caenophidia) from the Miocene of Gujarat, Western India: temporal constraints on dispersal of a derived snake: Journal of Vertebrate Paleontology, v. 27, p. 720–723.

Hunt, G., and Roy, K. 2006, Climate change, body size evolution, and Cope’s Rule in deepsea ostracodes. Proceedings of the National Academy of Sciences, v. 103, p. 1347–1352.

Hunt, G., 2008, Gradual or pulsed evolution: when should punctuational explanations be preferred?. Paleobiology, v. 34, p. 360–377.

Iqbal, M. W. A., 1980, Oligo-Miocene Bivalves & Gastropods from Kirthar Province, Lower Indus Basin, Pakistan: Geological Survey of Pakistan, v. 51, p. 1–59.

Jablonski, D., 1997, Body-size evolution in Cretaceous molluscs and the status of Cope’s rule: Nature, v. 385, p. 250–252.

Keeling, R. F., KöRtzinger, A., and Gruber, N., 2010, Ocean deoxygenation in a warming world: Annual Review of Marine Science, v. 2, p. 199–229.

Khaliq, I., Fritz, S. A., Prinzinger, R., Pfenninger, M., Böhning-Gaese, K., and Hof, C., 2015, Global variation in thermal physiology of birds and mammals: evidence for phylogenetic niche conservatism only in the tropics: Journal of Biogeography, v. 42, p. 2187–2196.

Kosnik, M. A., Jablonski, D., Lockwood, R., and Novack-Gottshall, P. M., 2006. Quantifying molluscan body size in evolutionary and ecological analyses: maximizing the return on data-collection efforts: Palaios, v. 21(6), p. 588–597.

Kroon D., Steens T., and Troelstra, S. R., 1991, Onset of monsoonal related upwelling in the western Arabian sea as revealed by planktonic foraminifers 1: Proceedings of the Ocean Drilling Program, Scientific Results, v. 1, p. 257–263.

Kulkarni, K.G., Bhattacharjee, S., and Borkar, V.D., 2007, Entobian bioerosion of Miocene oysters, Kachchh, Gujarat: Journal of the Geological Society of India, v. 69, p. 827–833.

Kulkarni, K.G., Bhattacharjee-Kapoor, S. and Borkar, V.D., 2010, Molluscan fauna from the Miocene sediments of Kachchh, Gujarat, India – Part 3. Gastropods: Journal of Earth System Science, v. 119, p. 307–341.

Kumar, P., Saraswati, P.K. and Banerjee, S., 2009, Early Miocene Shell Concentration in the Mixed Carbonate-Siliciclastic System of Kutch and their Distribution in Sequence Stratigraphic Framework: Journal of the Geological Society of India, v. 74, p. 432–444.

Less, G., Frijia, G., özcan, E., Saraswati, P. K., Parente, M., and Kumar, P., 2018, Nummulitids, lepidocyclinids and Sr-isotope data from the Oligocene of Kutch (western India) with chronostratigraphic and paleobiogeographic evaluations: Geodinamica Acta, v. 30, p. 183–211.

Mandic, O. and Piller, W.E., 2001, Pectinid coquinas and their palaeoenvironmental implications examples from the early Miocene of northeastern Egypt: Palaeogeography, Palaeoclimatology, Palaeoecology, v. 172, p. 171–191.

Mandic, O. and Steininger, F.F., 2003, Computer-based mollusc stratigraphy - a case study from the Eggenburgian (Lower Miocene) type region (NE Austria): Palaeogeography, Palaeoclimatology, Palaeoecology, v. 197, p. 263–291.

Mccall, J., Rosen, B., and Darrell, J., 1994, Carbonate deposition in accretionary prism settings: Early Miocene coral limestones and corals of the Makran Mountain Range in southern Iran: Facies, v. 31, p. 141 –178.

Mishra, V.P., 1976, Geology and Vertebrate Palaeontology of the Tertiary sequences of Western Kutch, Gujarat: Unpublished PhD. Thesis Lucknow University, p. 375.

Ohlberger, J., Mehner, T., Staaks, G., and Hölker, F., 2012, Intraspecific temperature dependence of the scaling of metabolic rate with body mass in fishes and its ecological implications: Oikos, v. 121, p. 245–251.

Patel, S. J., 2014, Ichnology of shallow marine argillaceous limestone, Miocene, Western Kachchh, India: Palaeontological Society of India Special Publication, p. 227–246.

Parmesan, C., 2006, Ecological and evolutionary responses to recent climate change: Annual Review of Ecology, Evolution, and Systematics, v. 37, p. 637–669.

Patnaik, R., Sharma, K.M., Mohan, L. and Blythe, A., 2014, Additional Vertebrate remains from the early Miocene of Kutch, Gujarat: Special Publication of the Palaeontological Society of India, v. 5, p. 335–351.

Poli, J.X., 1795, Testacea Utriusque Siciliae eorumque historia et anatome: Parma, Regio Typographeio, v. 2, p. 75–264.

Popov, S.V., Rögl, F., Rozanov, A.Y., Steininger, F.F., Shcherba, I.G., and Kovac, M., 2004, Lithological-Paleogeographic maps of Paratethys-10 maps Late Eocene to Pliocene: Courier Forschungsinstitut Senckenberg, v. 250, p. 1–46.

Raines, B.K. and Poppe, G.T., 2006, The family Pectinidae. A conchological iconography. ConchBooks, Hackenheim, Germany, pp. 720.

Renema, W., Bellwood, D.R., Braga, J.C., Bromfield, K., Hall, R., Johnson, K.G., and O’Dea, A., 2008, Hopping hotspots: global shifts in marine biodiversity: Science, v. 321, p. 654–657.

Reuter, M., Piller, W.E., and Erhart, C., 2012, A Middle Miocene carbonate platform under silici-volcaniclastic sedimentation stress (Leitha Limestone, Styrian Basin, Austria) — Depositional environments, sedimentary evolution and palaeoecology: Palaeogeography, Palaeoclimatology, Palaeoecology, v. 350–352, p. 198–211.

Reuter, M., Piller, W. E., Harzhauser, M., and Kroh, A., 2013, Cyclone trends constrain monsoon variability during Late Oligocene sea level highstands (Kachchh Basin, NW India): Climate Past, v. 9, p. 1–15.

RÖgl, V.F., 1998, Palaeogeographic Considerations for Mediterranean and Paratethys Seaways (Oligocene to Miocene): Naturhistorisches Museum Wien, v. 99, p. 279–310.

RÖGl, V.F., 1999, Short note Mediterranean and Paratethys. Facts and hypotheses of an Oligocene to Miocene Paleogeography (short overview): Geologica Carpathica, v. 50, p. 339–349.

Romdal, T. S., Colwell, R. K., and Rahbek, C., 2005, The influence of band sum area, domain extent, and range sizes on the latitudinal mid-domain effect: Ecology, v. 86, p. 235–244.

Roy, K., Jablonsk, D., and Valentine, J. W., 1995, Thermally anomalous assemblages revisited: patterns in the extraprovincial latitudinal range shifts of Pleistocene marine mollusks: Geology, v. 23, p. 1071–1074.

Sahni, A., and Mishra, V.P., 1975, Lower Tertiary Vertebrates from Western India: Monograph of the Palaeontological Society of India, v. 3, p. 1–48.

Sahni, A., and Mitra, H.C., 1980a, Neogene palaeobiogeography of the Indian subcontinent with special reference to fossil vertebrates: Palaeogeography, Palaeoclimatology, Palaeoecology, v. 31, p. 39–62.

Sahni, A. and Mitra, H.C., 1980b, Lower Miocene (Aquitanian-Burdigalian) palaeobiogeography of the Indian subcontinent: Geologische Rundschau, v. 69, p. 824–848.

Sarkar, D., Bhattacherjee, M., Chattopadhyay, D., 2019, Influence of regional environment in guiding the spatial distribution of marine bivalves along the Indian coast. Journal of the Marine Biological Association of the United Kingdom. v. 99, p. 163–177.

Scavia, D., Field, J. C., Boesch, D. F., Buddemeier, R. W., Burkett, V., Cayan, D. R., Fogarty, M., Harwell, M.A., Howarth, R.W., Mason, C., and Reed, D. J. (2002). Climate change impacts on US coastal and marine ecosystems: Estuaries, v. 25, p. 149–164.

şengör, A.M.C., 1981, The evolution of Palaeo-Tethys in the Tibetian segment of the Albides: in Proceeding Symposium Qinghai-Xizang (Tibet) Plateau: Science Press, Beijing, v. 1, p. 51–56.

SengöR, A. M. C., & Hsü, K. J. (1984). The Cimmerides of eastern Asia: history of the eastern end of Paleo-Tethys. Mémoires de la Société Géologique de France, v. 47, p. 139–167.

Smith, F. A., Betancourt, J. L., and Brown, J. H., 1995, Evolution of body size in the woodrat over the past 25,000 years of climate change: Science, v. 270, p. 2012–2014.

Sowerby, J. De C., 1840, Description of fossils from the Upper Secondary Formation of Cutch collected by C. W. Grant: Transactions of the Geological Society of London, v. 2, p. 327–329.

Sun, X., and Wang, P., 2005, How old is the Asian monsoon system?— Palaeobotanical records from China: Palaeogeography, Palaeoclimatology, Palaeoecology, v. 222, p. 181–222.

Tada, R., Zheng, H., and Clift, P. D., 2016, Evolution and variability of the Asian monsoon and its potential linkage with uplift of the Himalaya and Tibetan Plateau, Progress in Earth and Planetary Science, v. 3, p. 4.

Tewari, B.S., 1959, On a few fossil shark teeth from the Miocene Bed of Kutch, Western India: Proceeding of National Institute of Science India, v. 25, p. 230–236.

Thewissen, J.G.M., and Bajpai, S., 2008, New Oligocene mustelid from western India: Journal of Vertebrate Paleontology, v. 28, p. 565–567.

Thewissen, J.G.M. and Bajpai, S., 2009, New Skeletal Material of Andrewsiphius and Kutchicetus, Two Eocene Cetaceans from India: Journal of Paleontology, v. 83, p. 635–663. Unpublished Ph.D. Thesis, University Vienna.

Valentine, J. W. (1961). Paleoecologic molluscan geography of the Californian Pleistocene: University of California Publications in Geological Sciences, v. 34, p. 1–7.

Vitousek, P. M., Mooney, H. A., Lubchenco, J., and Melillo, J. M. (1997). Human domination of Earth’s ecosystems: Science, v. 277(5325), p. 494–499.

Vredenburg, E., 1925, Description of Mollusca from the post-Eocene Tertiary formations of north-western India: Cephalopoda, Opisthobranchiata, Siphonostomata: Memoirs of the Geological Survey of India, v. 50, p. 1–350.

Vredenburg, E., 1928, Description of Mollusca from the post-Eocene Tertiary formations of north-western India. 2: Memoirs of the Geological Survey of India, v. 50, p. 351–463.

Zhuang, G., Pagani, M., & Zhang, Y. G. (2017). Monsoonal upwelling in the western Arabian Sea since the middle Miocene: Geology, v. 45, p. 655–658.

Zunino, M. and Pavia, G., 2009, Lower to Middle Miocene Mollusc assemblages from the Torino Hills (NW Italy): Synthesis of new data and chronostratigraphical arrangement: Rivista Italiana di Paleontologia e Stratigrafia, v. 115, p. 349–370.

